# *In-vivo* crystals reveal protein interactions

**DOI:** 10.1101/778779

**Authors:** Eleanor R. Martin, Alessandro Barbieri, Robert C. Ford, Robert C. Robinson

## Abstract

Crystallisation of recombinant proteins has been fundamental to our understanding of protein function, dysfunction, and molecular recognition. However, this information has often been gleaned under non-physiological extremes of protein, salt, and H^+^ concentrations. Here, we describe the development of the robust iBox-PAK4cat system that spontaneously crystallises in several mammalian cell types. The developments described here allow the quantitation of *in-vivo* protein-protein interactions using a novel GFP-linked reporter system. Here, we have combined this assay with *in-vitro* X-ray crystallography and molecular dynamics studies characterise the molecular determinants of the interaction between NHERF1 PDZ2 and CFTR, a protein complex pertinent to the genetic disease cystic fibrosis. These studies have revealed the crystal structure of the extended PDZ domain of NHERF1, and indicated, contrary to previous reports, that residue selection at −1 and −3 PDZ-binding motif positions influence the affinity and specificity of the interaction. The results presented here demonstrate that the iBox-PAK4cat assay could easily be utilised to screen other protein-protein interactions.

## Introduction

The catalytic domain of the serine/threonine kinase PAK4 (PAK4cat) and its endogenous inhibitor Inka1 have recently been shown to readily form rod-type crystals on co-transfection into a variety of mammalian cell types [1]. Crystals formed following transfection of a fusion construct comprising of a central 38 residue region of Inka1, termed the Inka1-Box (iBox), and PAK4cat (iBox-PAK4cat) were diffracted at the synchrotron beamline, enabling the first *in-cellulo* human protein structure to be determined to 2.95 Å (PDB: 4XBR). The crystal lattice revealed that iBox-PAK4cat forms a hexagonal array with channels 80 Å in diameter that run the length of the crystal. The size of this cavity allowed a guest protein (GFP) be incorporated into the crystal lattice [1]. This, as well as the relative ease with which crystals can be generated with following transfection of DNA into mammalian cells, suggests that iBox-PAK4cat crystals may be a useful tool for variety of experimental purposes, including crystallisation of guest proteins, use as environmentally responsive cellular sensors, and use in protein-protein interaction (PPI) screening assays. The potential uses of iBox-PAK4cat *in-cellulo* crystals however, for both structural and other applications, are relatively unexplored and require further study.

PSD-95/Dlg-1/ZO-1 (PDZ) domains constitute one of the most commonly identified protein interaction modules, with over 500 PDZ domain-containing proteins identified in the human genome alone [2]. PDZ domains characterized to date share a common core structure, consisting of a 6-stranded antiparallel beta sandwich structure (β1-β6) flanked by an alpha helix on each side (α1, α2). PDZ domains fulfil their scaffolding function through the recognition of short linear PDZ-binding motifs which are commonly found at the extreme C-termini of receptors and ion channels. PDZ-binding motifs are generally agreed to comprise of four residues, referred to by convention as P0, P-1, P-2, and P-3 from the C-terminal residue inward. Early studies characterising PDZ domain-motif interactions led to the classification of PDZ domains based on their preferences for particular PDZ-binding motifs, indicating motif selection is primarily dictated by the motif residues at P0 and P-2. Peptide library screening studies identified PDZ domains which displayed distinct preferences for the type I motif X-[S/T]-X-Φ (where Φ is a hydrophobic amino acid and X is any amino acid), the type II motif X-Φ-X-Φ [3, 4], or the less common type III motif X-[D/E]-X-Φ [5].

The cystic fibrosis transmembrane conductance regulator (CFTR), or ABCC7, is a chloride ion channel expressed in the epithelia of the airways, intestine, pancreas, sweat gland, and testis [6]. Loss-of-function mutations of CFTR cause the autosomal-recessive disease cystic fibrosis (CF). CF is the most prevalent genetic disease amongst European populations, affecting on average 1 in every 3500 live births [7]. The most common CF-causing mutation is the deletion of phenylalanine at position 508 (F508del), which is present on at least one allele in 90 % of patients in some populations [8]. The F508del mutation causes protein-misfolding and subsequent targeting of CFTR for degradation via the endoplasmic reticulum-associated degradation (ERAD) pathway [9]. CFTR contains the conserved type-I PDZ-binding motif (D/E)T(R/K)L at its C-terminus. Through this motif, CFTR is known to interact with several PDZ domain-containing proteins, including CFTR-associated ligand (CAL), Shank2, as well as all four members of the NHERF family (NHERF1, NHERF2, NHERF3, and NHERF4) [10–13]. NHERF1 is comprised of two PDZ domains (PDZ1 and PDZ2), both of which are able to bind to the PDZ-binding motif of CFTR [10, 14–17]. NHERF1 has been shown to increase both the activity and plasma membrane localisation of both wild-type and F508del CFTR [18–23]. While mutations of the CFTR C-terminus are rare, they have been reported as being disease-causing [24]. Furthermore, CFTR variants in which the CFTR PDZ-binding motif is compromised through either truncation or substitution (e.g. S1455X, Q1476X, L1480P) have been associated with various CF-related disorders such as elevated sweat chloride concentration and congenital bilateral absence of the vas deferens (see http://www.genet.sickkids.on.ca). It has been demonstrated that while F508del CFTR interacts poorly with NHERF1, the CFTR corrector VX-809 increases the binding affinity between these two proteins [25]. Together, these studies suggest that one potential strategy for the further development of CF therapeutics may be the generation of small molecule corrector-type drugs specifically targeted at enhancing the interaction between NHERF1 and CFTR. Indeed, more generally, there is significant interest in targeting specific PDZ domain-motif interactions with small molecule inhibitors or peptides which block the PDZ binding site [26, 27]. The development of drugs which target certain PDZ domains but not others however remains challenging and requires further characterisation of the molecular determinants of specific PDZ domain-motif interactions.

The experiments described here employ a variety of approaches to further characterise the molecular determinants of the interaction between the second PDZ domain of NHERF1 and the CFTR. These results indicate that the PDZ-binding motif of CFTR appears to be an optimal PDZ-binding motif for interaction with PDZ2 of NHERF1, and that residue selection at the −1 and −3 motif positions appears to influence the affinity and specificity of the interaction. These findings have been corroborated *in-cellulo* through the development of a novel PPI screening assay based on the iBox-PAK4cat crystallisation system. This study demonstrates that iBox-PAK4cat crystals represent a potentially useful tool for the study of PDZ domain-motif interactions in a cellular context and may be a useful tool in future screening studies.

## Results

### The structure of the extended NHERF1 PDZ2 domain

To understand fully the determinants of the NHERF1 PDZ2-CFTR interaction we sought to determine the co-crystal structure. In order to facilitate co-crystallisation, a chimeric construct was generated in which the NHERF1 PDZ2 domain (NHERF1 residues 150-270) was fused to a C-terminal DTRL tetrapeptide motif corresponding to the last four residues of CFTR (CFTR residues 1477-1480), mirroring an approach commonly used for crystallisation of PDZ domain-motif complexes [28–31]. This strategy produced diffraction-quality crystals, enabling the structure to be solved and refined to 2.2 Å (Table 1).

**Table 1:** Data collection and refinement statistics for the NHERF1 PDZ2-CFTR complex

| **Data collection** |  |
| --- | --- |
| Space group | I2<sub>1</sub>2<sub>1</sub>2<sub>1</sub> |
| Cell dimensions |  |
| (a, b, c) (Å) | 84.9, 85.0, 89.6 |
| (α, β, γ) (°) | 90.0, 90.0, 90.0 |
| Resolution (Å) | 30.1-2.20 (2.28-2.20) |
| R<sub>merge</sub> (%) | 17.6 (95.1) |
| R<sub>pim</sub> (%) | 11.3 (60.0) |
| CC<sub>1/2</sub> | 0.984 (0.634) |
| I/σ(I) | 4.3 (1.6) |
| Unique reflections | 16793 (1653) |
| Completeness (%) | 99.3 (97.8) |
| Multiplicity | 3.5 (3.6) |
| **Refinement** |  |
| Resolution (Å) | 19.8-2.20 (2.28-2.20) |
| No. of reflections: working/test | 16732/780 (1618/75) |
| R<sub>work</sub>/R<sub>free</sub> | 23.6/27.5 (33.7/35.7) |
| No. of atoms | 2017 |
| Protein | 1926 |
| Water | 91 |
| Average B-factor (Å<sup>2</sup>) | 39.9 |
| Protein (Å<sup>2</sup>) | 39.9 |
| Water (Å<sup>2</sup>) | 39.0 |
| RMS deviations |  |
| Bond lengths (Å) | 0.009 |
| Bond angles (Å) | 1.36 |
| Ramachandran |  |
| Favoured (%) | 98.8 |
| Allowed (%) | 1.2 |
| Disallowed (%) | 0.0 |

The overall topology of the canonical NHERF1 PDZ2 domain is consistent with standard PDZ domains, comprising of a 6-stranded β-barrel (β1-β6) and two α-helices (α1 and α2) (Fig. 1a). In agreement with previous reports [17], the structure revealed that NHERF1 PDZ2 contains additional structured extensions beyond the core PDZ-fold. These extended regions form an α-(α3) and a 3_10_ (α4) helix, which pack against the β1, β4, and β6 strands through the formation of a closed hydrophobic cluster (Fig. 1b). While this hydrophobic cluster is formed from residues distal to the binding site, the extended NHERF1 PDZ2 domain has been shown to have an almost 18-fold higher binding affinity for the CFTR C-terminus than the canonical domain [17]. Comparison of the structure of the extended NHERF1 PDZ2 with the canonical domain (NHERF1 residues 148-239) in the apo form (PDB: 2OZF) reveals high structural similarity, with an RMSD of 0.74 Å for the 85 equivalent Cα atoms (Supplementary Fig. 1). This indicates that the extended region does not induce significant structural changes in the core PDZ domain, but rather conveys its effects through allosteric interactions which mediate the stability of the binding site. Interestingly, multiple sequence alignment and secondary structure predictions indicate that similar structured extensions outside of the core PDZ fold are common to several members of the NHERF family (Fig. 3b, Supplementary Fig. 2), although thus far only those of NHERF1 have been characterised.

**Figure 1:**
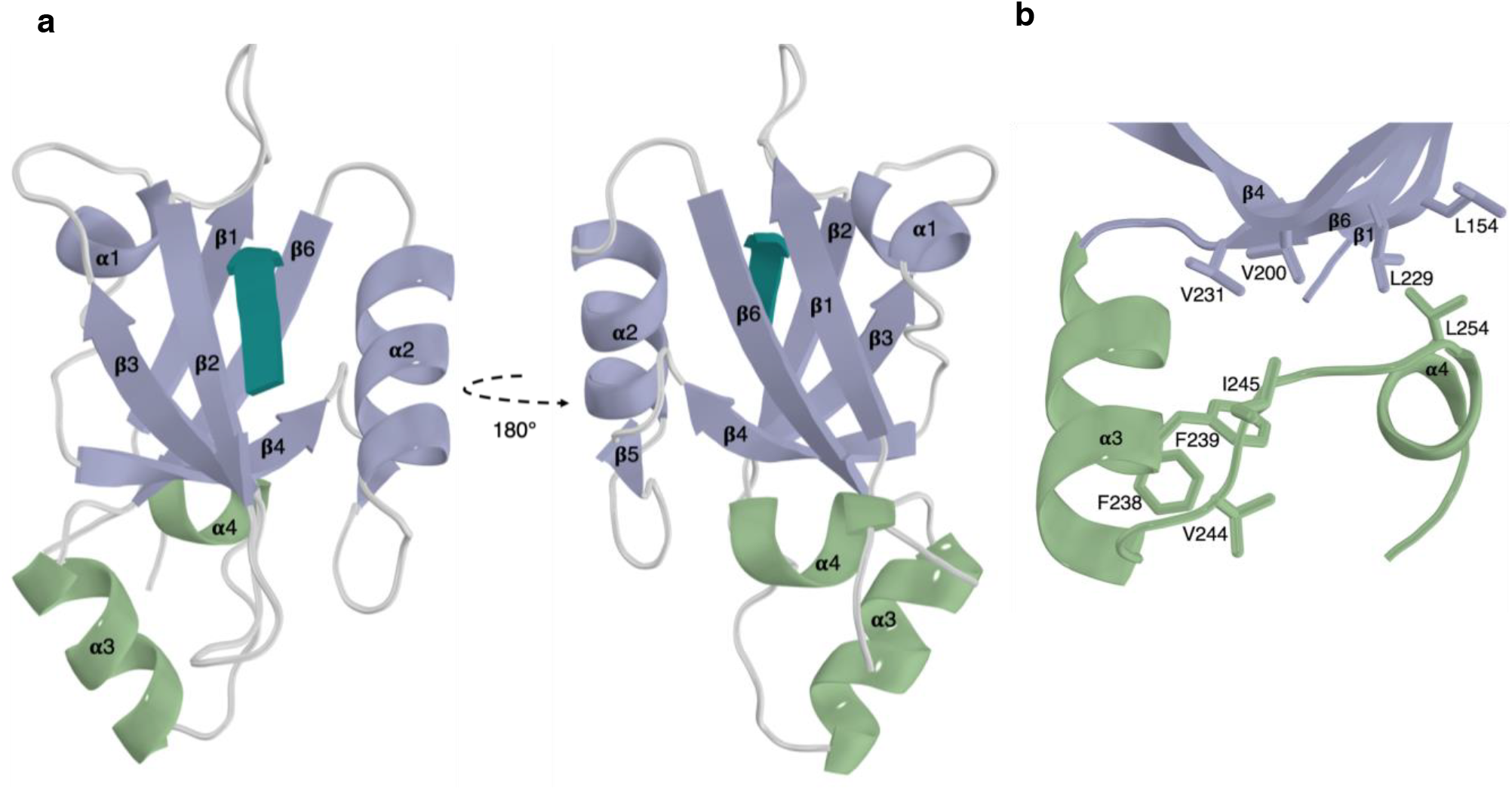
The extended PDZ2 domain of NHERF1 in complex with the CFTR PDZ-binding motif. **(a)** The extended helical regions (green) couple to the canonical PDZ domain (purple) at a region distal to the PDZ-binding motif (teal). **(b)** Hydrophobic interactions formed between the α3 and α4 and β1, β4, and β6 from the core PDZ fold.

### The determinants of the NHERF1 PDZ2-CFTR interaction

The PDZ binding motif of CFTR inserts into the β2-α2 binding groove in a canonical fashion (Fig. 1a, 2a), extending the β-sheet of the PDZ domain and burying a solvent accessible surface area of 876 Å^2^. The DTRL peptide at the CFTR C-terminus represents a typical type I PDZ-binding motif, and as such, the structure presented here shares many common features with a typical type I PDZ domain-motif interaction. The T-2 residue of the PDZ binding motif can be observed to form a hydrogen bond with a conserved histidine residue at position 212 of NHERF1 PDZ2 (Fig. 2a), consistent with a shared mechanism of recognition found in type I interactions. The L0 carboxyl group and leucine side chain appear to be key determinants of binding through interactions with the conserved ^163^GYGF^167^ carboxylate-binding motif and hydrophobic binding pocket respectively (Fig. 2). In particular, the binding pocket appears to be stereochemically adapted to accommodate the leucine side chain (Fig. 2b). Other residues of the PDZ-binding motif can be observed to take part in numerous interactions with the NHERF1 PDZ2 domain. Specifically, the D-3 side chain forms a salt bridge with R180 and a hydrogen bond with H169, and R-1 forms a salt bridge with D183 and a hydrogen bond with N167 (Fig. 2a).

**Figure 2:**
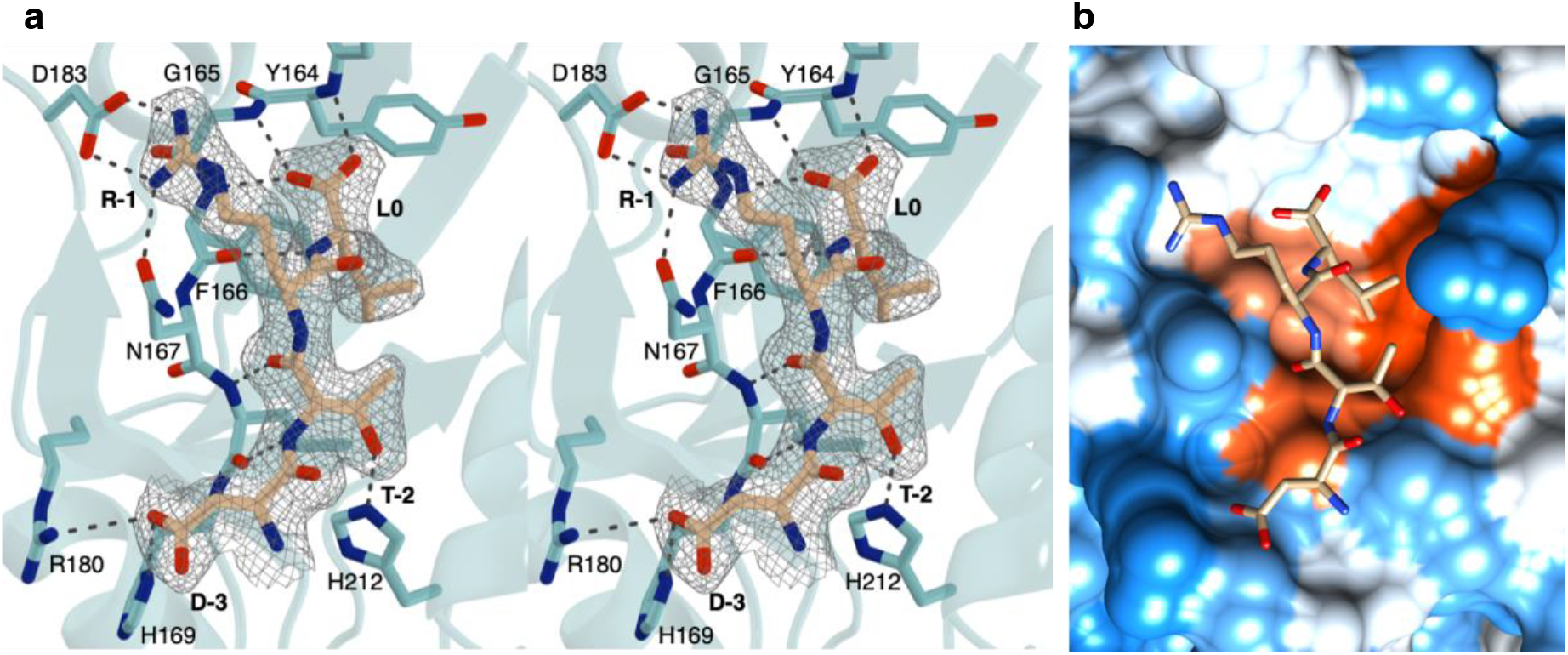
Interactions between NHERF1 PDZ2 and CFTR. **(a)** Stereo view of the NHERF1 PDZ2 binding site (cyan) bound to the PDZ-binding motif of CFTR (beige). The 2Fo-Fc map is shown around the CFTR PDZ-binding motif contoured at 1.0 sigma within 1.5 Å of the selected atoms. **(b)** Hydrophobicity surface structure of NHERF1 PDZ2 binding site based on the Kyte-Doolittle scale with the most hydrophillic regions shown in blue and most hydrophobic regions shown in orange-red. The CFTR PDZ-binding motif is shown as a stick model in beige. Image was generated using Chimera [32].

Several studies have indicated that the CFTR C-terminus binds to NHERF1 PDZ2 with lower affinity than PDZ1 [10, 17, 33]. Indeed, while NHERF1 PDZ1 and PDZ2 domains share 54.4 % sequence identity, they have been shown to display distinct differences with regards to ligand binding selectivity and affinity. The PDZ domains of NHERF1 are highly structurally similar (Fig. 3a). However, natural sequence variation of a number of key residues within the binding pocket can be observed, including the substitution of E43 within PDZ1 with an aspartate residue at the equivalent position (D183) within PDZ2 (Fig. 3b). Studies have suggested that the lower binding affinity of PDZ2 can be attributed to the shorter side-chain of D183 that cannot form a salt-bridge with R-1 residue of the CFTR PDZ-binding motif [17, 34]. The crystal structure presented here contradicts this hypothesis, since the salt-bridge between D183 and R-1 is present (Fig. 2a, 3a). The majority of known CFTR-binding PDZ domains contain a negatively charged residue at this position within the binding pocket, including those outside of the NHERF family such as Shank1 (Fig. 3b), implying that it is charge rather than side-chain length that is of importance. Furthermore, a positively charged residue at P-1 of the PDZ-binding motif is conserved cross-species for CFTR (Supplementary Fig. 3). As such, these results indicate that the formation of an ionic interaction between these two residues may be a common mechanism for recognition and selectivity between CFTR and its respective CFTR-binding PDZ domains.

**Figure 3:**
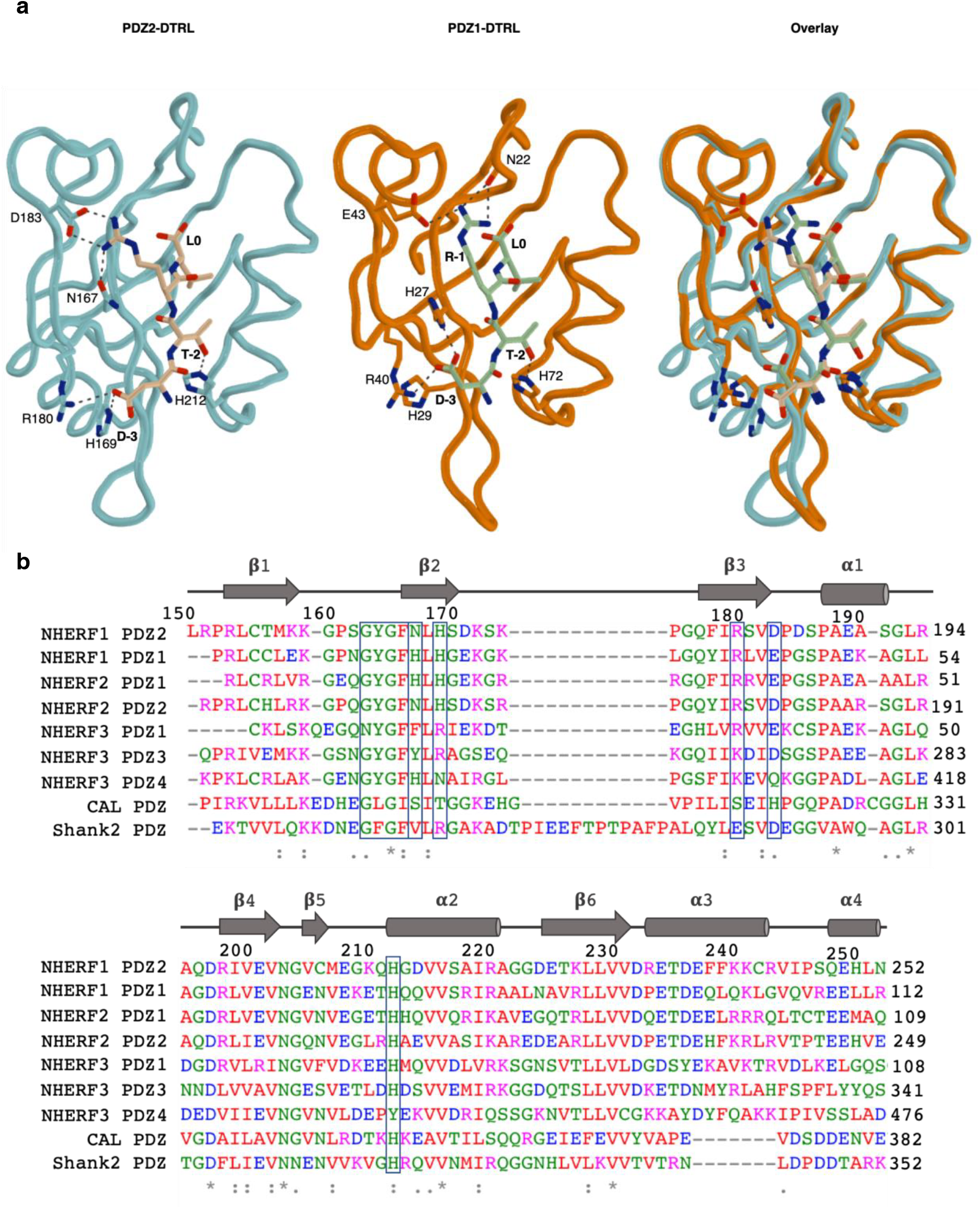
Comparison of human CFTR-binding PDZ domains. **(a)** NHERF1 PDZ2-DTRL, NHERF1 PDZ1-DTRL (PDB: 1I92), and overlay structures are shown as indicated. NHERF1 PDZ2-DTRL is shown as previously. NHERF1 PDZ1 is shown in orange and the PDZ-binding motif of CFTR in green. In both cases side-chain groups of residues relevant to the ligand interaction are indicated. Black dashes represent bond distances measuring 3.3 Å or less. **(b)** Multiple sequence alignment of known CFTR-binding PDZ domains. Residue numbers and secondary structure of NHERF1 PDZ2 are indicated above. Conserved ‘X-ϕ-G-ϕ’ carboxylate-binding loop and key residues within the PDZ binding pocket, including D183/E43, are boxed.

### Use of iBox-PAK4cat *in-cellulo* crystals as a tool for studying protein-protein interactions

The *in-cellulo* crystal structure of iBox-PAK4cat revealed a hexagonal lattice with channels ~80 Å in diameter that run the length of the crystal [1]. We hypothesised that it may be possible to introduce NHERF1 PDZ2 into the crystal lattice in order facilitate its *in-cellulo* crystallisation. A NHERF1 PDZ2 containing iBox-PAK4cat fusion construct (iBox-PDZ2-PAK4cat) readily formed diffraction-quality crystals (Supplementary Fig. 4, Supplementary Table 1). A partial electron density map was revealed for the PDZ2 domain, although it was of insufficient quality to trace the full polypeptide chain (Supplementary Fig. 5), indicating that additional construct optimisation is required to stabilise guest proteins in a single orientation within the crystal lattice. GFP-containing fluorescent crystals have also been produced through co-transfection of PAK4cat with various GFP-Inka1 constructs [1]. In an extension of these approaches, we postulated that, through monitoring of GFP fluorescence levels, iBox-PAK4cat crystals could also be utilised as high-density *in-vivo* sensors for the detection of protein-protein interactions. In particular, we sought to utilise the iBox-PAK4cat system as a tool to enable detection of PDZ domain-motif interactions. This approach is summarised in Figure 4a. Two crystal forming constructs were tested, one containing a C-terminal tetrapeptide corresponding to the CFTR PDZ-binding motif (iBox-PAK4cat-DTRL), and a negative control containing a non PDZ-binding C-terminal motif (iBox-PAK4cat-AAAA). While both iBox-PAK4cat-DTRL and iBox-PAK4cat-AAAA constructs appeared to be similar in terms of crystal generation proficiency (Supplementary Fig. 6), iBox-PAK4cat-DTRL crystals demonstrated clearly observable GFP fluorescence on co-transfection with GFP-NHERF1PDZ1. Co-transfection with iBox-PAK4cat-AAAA, however, resulted in only cytoplasmic GFP fluorescence, with minimal fluorescence observable in the crystals (Fig. 4b). The easily visible crystals formed from GFP–NHERF1PDZ and iBox-PAK4cat-DTRL co-transfection allowed for time-lapse analysis of crystal formation (Supplementary Movie 1).

**Figure 4:**
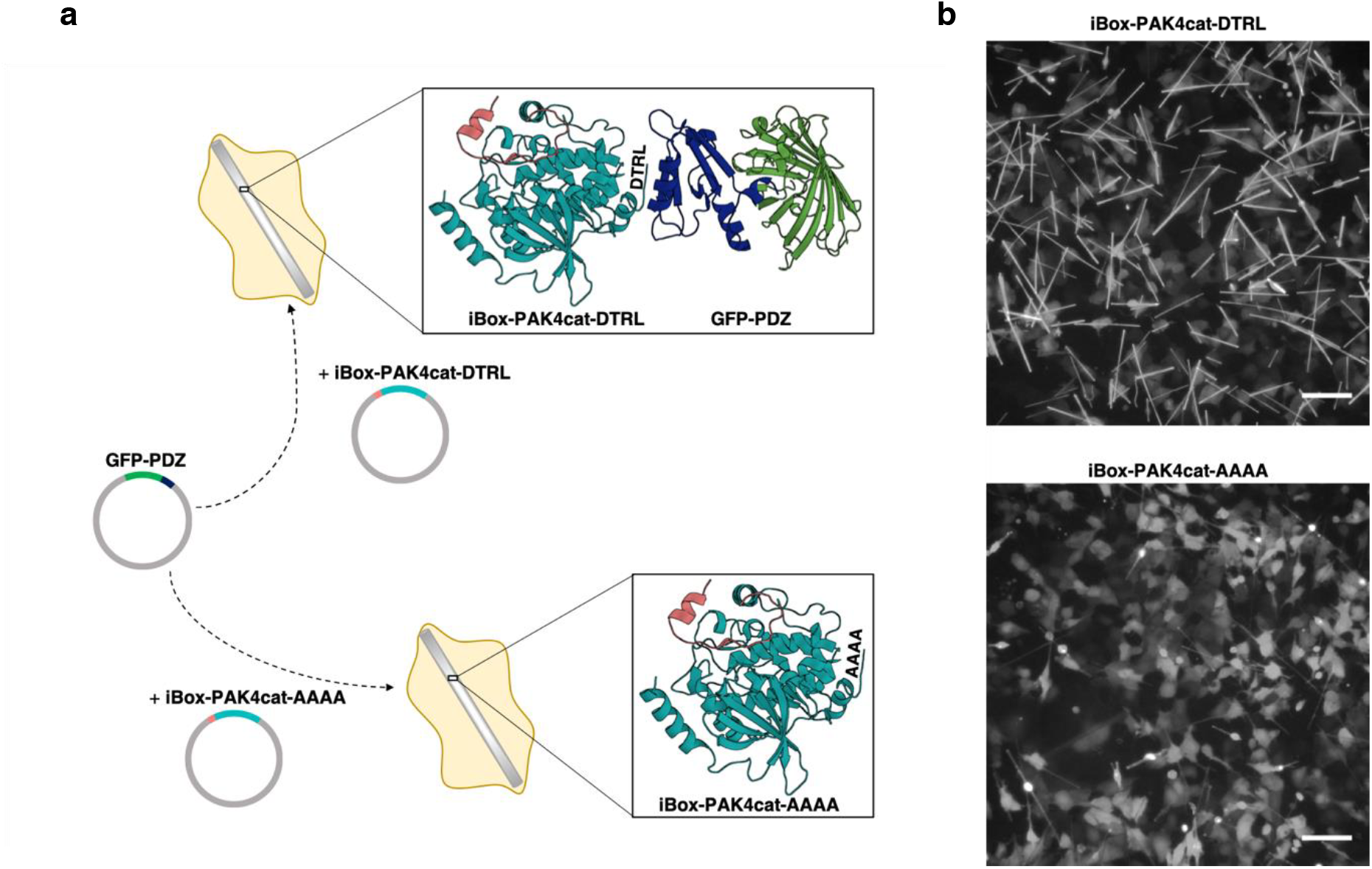
Use of iBox-PAK4cat *in-cellulo* crystals as a tool for studying protein-protein interactions. **(a)** Schematic of the iBox-PAK4cat PPI assay. Mammalian cells are co-transfected with GFP-PDZ and iBox-PAK4cat constructs containing a C-terminal tetrapeptide motif (iBox-PAK4cat-DTRL and iBox-PAK4cat-AAAA). For iBox-PAK4cat-AAAA no interaction takes place and GFP-PDZ is not incorporated into the iBox-PAK4cat crystal lattice. Conversely, for iBox-PAK4cat-DTRL a PDZ domain-motif interaction occurs and GFP-PDZ is incorporated into the iBox-PAK4cat crystal lattice. The iBox-PAK4cat (red and teal, respectively) *in-cellulo* crystal structure (PDB: 4XBR) bound to GFP-PDZ (green and blue, respectively) is modelled for illustrative purposes only. **(b)** Fluorescent images of cells co-transfected with GFP-NHERF1PDZ1 and iBox-PAK4cat-DTRL or iBox-PAK4cat-AAAA, as indicated. Equivalent transmitted light images are shown in Supplementary Fig. 6. Scale bar bottom right indicates 100 μm.

### iBox-PAK4cat *in-cellulo* protein-protein interaction assay reveals the determinants of NHERF1 PDZ domain binding selectivity

Since iBox-PAK4cat crystals enable clear visual differentiation between binding and non-binding events on co-transfection with GFP-PDZ (Fig. 4b), we sought to determine whether this system could also detect changes to the PDZ-binding motif. A number of constructs were generated in which residues of the CFTR PDZ-binding motif were mutated to alanine, as well as a phosphomimetic motif in which the threonine of the PDZ-binding motif was mutated to glutamic acid (Fig. 5a). On co-transfection with GFP-PDZ constructs, visual comparison indicated that these substitution mutants demonstrated lower levels of crystal fluorescence than the iBox-PAKcat-DTRL construct (Fig. 5b, Supplementary Fig. 7). In particular, introduction of the C-terminal leucine to alanine substitution mutant (iBox-PAK4cat-DTRA) had a drastic effect, with crystal fluorescence levels appearing similar to the iBox-PAK4cat-AAAA negative control (Fig. 5b, Supplementary Fig.7). That such a large effect comes from a relatively conservative amino acid change may be understood by the stereochemical adaptation of the binding pocket to leucine observed in the crystal structure (Fig. 2b).

**Figure 5:**
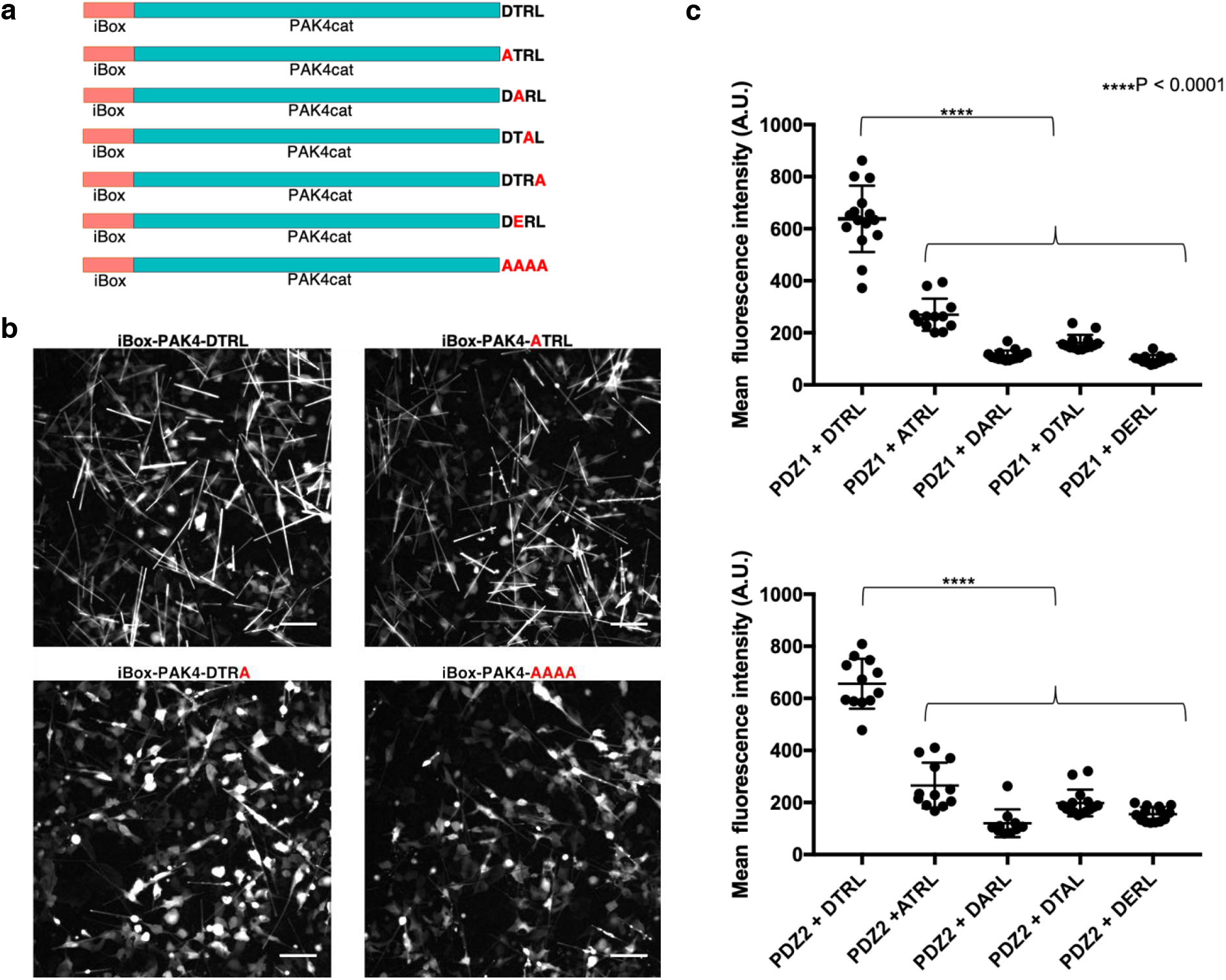
Effect of PDZ-binding motif mutations on GFP integration into iBox-PAK4cat crystals. **(a)** iBox-PAK4cat constructs utilised in this study. **(b)** Fluorescent images of cells co-transfected with GFP-NHERF1PDZ2 and iBox-PAK4cat constructs, as indicated. Equivalent transmitted light images are shown in Supplementary Fig. 8. Scale bar bottom right indicates 100 µm. (c) Substitution mutations of the DTRL PDZ-binding motif are associated with significantly lower levels of fluorescence within iBox-PAK4cat crystals (two-way ANOVA with Sidak’s multiple comparisons test, F_(4, 122)_ = 287, P < 0.0001). Each point represents the mean fluorescence intensity of a single image containing multiple crystals, as shown in Supplementary Fig. 9. N = 10-15.

To characterise these constructs further, we sought to quantify the intensity of GFP fluorescence within the crystals. To remove background fluorescence caused by GFP-PDZ expression in the surrounding cells, crystal-containing cells were lysed, and crystals were imaged in a cell-free form (Supplementary Fig. 9). For co-transfections employing iBox-PAK4cat-DTRA constructs levels of fluorescence were not above control levels. For the remaining constructs, quantitative analysis indicated that ATRL, DARL, DTAL, and DERL containing iBox-PAK4cat constructs demonstrated significantly lower levels of fluorescence than that of iBox-PAK4cat-DTRL (Fig. 5c). As such, it is evident that all residues of the CFTR ‘DTRL’ PDZ-binding motif form specific interactions within the binding pockets of NHERF1 PDZ1 and PDZ2.

### Molecular dynamics simulations and binding free energy studies on NHERF1 PDZ2-CFTR complex

The crystal structure presented here, supported by the findings of the novel iBox-PAK4cat *in-*cellulo PPI assay, indicate that side-chain mediated interactions within the NHERF1 PDZ2 binding pocket enable the specific recognition of the CFTR PDZ-binding motif. In order to explore these findings further, *in-silico*, molecular dynamics (MD) simulations were performed.

Following evaluation and verification of the protonation states of the histidine residues within the binding pocket (Materials and Methods, Supplementary Fig. 10), MD simulations were carried out using the obtained NHERF1 PDZ2-DTRL crystal structure. Simulations confirmed the persistence of hydrogen bonds observed in the crystal structure. R-1 was found to steadily form a salt-bridge with the carboxylate of D183, as the interactions between the carboxylic group of the D-3 with the guanidino-group of R180 and the side chain of H169. Similarly, the interaction between the hydroxyl group of T-2 and the side chain of H212 showed a clear occupancy for 83-85% of the 50 ns production run. Two different binding modes could be observed for the terminal carboxylate group of L0: the first, similar to the initial structure and involving direct interaction with the conserved GYGF carboxylate-binding motif; the second, with the terminal carboxylate shifted and more inclined towards R220 in the opposite direction (Fig. 6). Surprisingly, in this mode, the initial direct interactions between the L0 terminal carboxylate and the GYGF motif are mediated by a water molecule which steadily occupies the space vacated by the carboxylate during its shift towards R220. This water molecule forms a network of interactions involving L0 and the GYGF motif when there are no direct interactions among the residues. A shift in the R-1 side-chain could also be observed, enabling the interaction of the R-1 side-chain with the backbone carbonyl of S162 and a strong stabilization of the already present interaction with the amidic carbonyl group of the N167 side-chain. This orientation of the R-1 side-chain appears to be an intermediate between the those observed in the PDZ1 and PDZ2 crystal structures (Fig. 3a), and is coupled with L0 carboxylate shift and tightening of the β1-β2 loop (Fig. 6).

**Figure 6:**
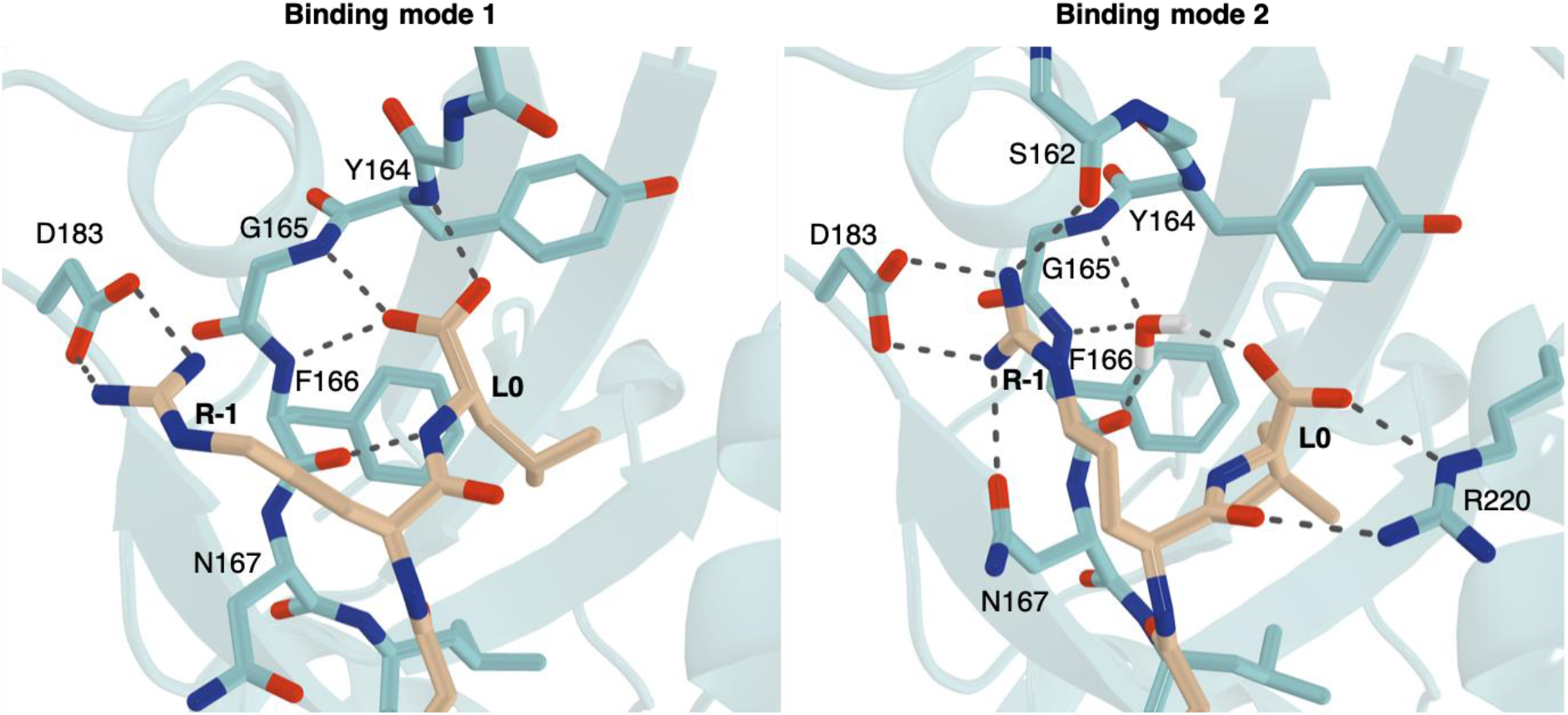
Binding modes observed for L0 and R-1 in molecular dynamics simulations. The two binding modes shown correspond to a 50:50 distribution between the two states. Representative frames of two binding modes were selected after clustering of two MD simulations. NHERF1 PDZ2 is coloured in cyan and the CFTR PDZ-binding motif in beige. Black dashes represent bond distances measuring 3.3 Å or less. For clarity, hydrogens are shown for the water molecule only.

In order to confirm the findings of the iBox-PAK4cat protein-protein interaction assay, the same protocol was utilized to perform MD simulations on NHERF1 PDZ2 in complex with the ATRL, DARL, DTAL, and DTRA alanine substitution mutants of the PDZ-binding motif. For these simulations, the RMSD values related to the secondary structure elements were found to remain in the same range as for the WT complex. However, the stability of the peptide for all alanine substitution mutants was found to be much lower (Supplementary Fig. 11), presumably due to the absence of key interactions within the binding pocket. In order to determine the effect of these mutations on interaction affinity, binding free energies for both replicates were calculated using both the Solvation Interaction Energies (SIE) and Molecular Mechanics Poisson–Boltzmann Surface Area (MM-PBSA) methods (Table 2). During the initial evaluation of different protonation states on the wild type simulations, the estimated binding energies were all found to be in agreement using both methods. However, with the MM-PBSA method only the ATRL and the DTAL systems were found to be substantially less favourable in terms of binding energies, while using the SIE method, and in agreement with the findings of the iBox-PAK4cat PPI assay, all alanine substitution mutants were found to be less energetically favourable than the WT motif. In particular, the presence of the guanidino group at P-1 appears to be critical to the formation of interactions within the binding pocket, with a decrease over 4 kcal/mol (SIE method) and 15 kcal/mol (MM-PBSA) observable for both replicas of the DTAL system.

**Table 2.** Binding Free Energies for WT and Alanine mutants with MMPBSA and SIE methods.

| | $\Delta G$ kcal/mol<br>(± SEM)<br>DTRL | $\Delta G$ kcal/mol<br>(± SEM) ATRL | $\Delta G$ kcal/mol<br>(± SEM)<br>DARL | $\Delta G$ kcal/mol<br>(± SEM)<br>DTAL | $\Delta G$ kcal/mol<br>(± SEM)<br>DTRA |
| --- | --- | --- | --- | --- | --- |
| **MM-PBSA** | -73.24 ± 0.37<br>-75.51 ± 0.40 | -63.17 ± 0.34<br>-64.55 ± 0.30 | -72.66 ± 0.35<br>-74.72 ± 0.38 | -51.37 ± 0.52<br>-58.28 ± 0.33 | -72.46 ± 0.40<br>-73.69 ± 0.34 |
| **SIE** | -14.53 ± 0.05<br>-15.08 ± 0.03 | -10.88 ± 0.03<br>-10.93 ± 0.02 | -11.33 ± 0.03<br>-11.51 ± 0.03 | -10.46 ± 0.04<br>-10.36 ± 0.02 | -11.27 ± 0.03<br>-11.43 ± 0.03 |

## Discussion

The findings presented here corroborate published data, which indicate that NHERF1 PDZ2 contains structured extensions outside of the core PDZ fold [16, 17]. Structured extensions outside of the canonical PDZ domain are not unique to NHERF1 PDZ2. NHERF1 PDZ1 has been shown to contain a highly similar C-terminal helical region, albeit with only comparatively negligible effects on stability and ligand binding affinity [17]. The results presented here indicate that similar structured extensions are common to several members of the NHERF family (Fig. 3b, Supplementary Fig. 2). Indeed, bioinformatics studies have suggested that 40 % of PDZ domains contain structured extensions outside of the core PDZ fold [35]. One of the best characterised examples to date is the third PDZ domain of PSD-95, which also contains a C-terminal α3 helix. The overall characteristics of this C-terminal extension are very similar to the extended region of NHERF1 PDZ2 in that i) the extension is distal to the binding site, ii) the extension does not induce any major structural changes, and iii) the extension increases binding affinity for the ligand 21-fold [36]. As such, extension mediated allosteric stabilization of the binding site may be a common mechanism shared among several PDZ domains. Further structural and biochemical studies are required in order to further elucidate the role of these structured extensions across the NHERF family.

This study has also provided further insight into the determinants of interaction between NHERF1 PDZ2 and the CFTR PDZ-binding motif. While previous studies had suggested that the E43 to D183 substitution in PDZ2 was the sole determinant for the reduced binding affinity of NHERF1 PDZ2 to the CFTR C-terminus relative to PDZ1 [17, 34], the crystal structure presented here demonstrates that D183 is able to maintain the interaction with R-1 (Fig. 2a). These findings were confirmed by MD simulations, where this salt-bridge interaction is found to be steady for the entire production (Fig. 6), indicating that the higher binding affinity achieved by NHERF1 PDZ1 is likely to be a result of a variety structural features throughout the binding pocket, rather than sequence variation of a single residue. Interestingly, the MD simulations indicated two distinct binding modes for

R-1 and L0 within the binding pocket. In addition to the canonical interaction between L0 and the carboxylate-binding loop, L0 can also be observed to shift towards R220, with a water molecule instead facilitating the indirect interaction between the carboxylate group and the GYGF motif. Water-mediated hydrogen bonds between the terminal residue of the binding motif and the PDZ binding pocket have been reported previously [28, 37], although the mode observed here appears to be mechanistically distinct. This orientation of L0 is also associated with the formation of a hydrogen bond between the R-1 side-chain with the backbone carbonyl of S162, a finding which is in-keeping with previous reports that phosphorylation of S162 disrupts the interaction between NHERF1 PDZ2 and the PDZ-binding motif of CFTR [15].

The novel iBox-PAK4cat protein-protein interaction assay presented here has provided further insight into the determinants of the interaction between NHERF1 and CFTR. This approach allows clear differentiation between PDZ domain binding (iBox-PAK4cat-DTRL) and non-binding (iBox-PAK4cat-AAAA) motifs through visual comparison of crystal fluorescence (Fig. 4b). Introduction of the C-terminal leucine to alanine substitution mutant (iBox-PAK4cat-DTRA) had a drastic effect on GFP integration into the crystals, with crystal fluorescence levels appearing similar to the iBox-PAK4cat-AAAA negative control (Fig. 5b). Such a finding is supported by co-immunoprecipitation data [38] and is consistent with the finding that the binding pocket is stereochemically adapted for the accommodation of the leucine side-chain (Fig. 2b). As such, it is likely that the much smaller side-chain of alanine would produce a much less energetically favourable interaction. The phosphomimetic motif-containing iBox-PAK4cat-DERL construct was also found to result in reduced fluorescence levels (Fig. 5c). Phosphorylation of the serine or threonine residue at this position has been demonstrated to disrupt the PDZ domain-motif interaction in several other cases [39–41]. While phosphorylation of this residue in CFTR has been hypothesised previously [15], the role of phosphorylation as a regulatory mechanism for this interaction requires further investigation.

Quantitative analysis of iBox-PAK4Cat-ATRL, iBox-PAK4Cat-DARL, and iBox-PAK4cat-DTAL constructs indicated that these PDZ-binding motif mutants had significantly lower levels of GFP integration into iBox-PAK4cat crystals than the wild-type CFTR PDZ-binding motif (Fig. 5c). This supports the finding, as demonstrated by the crystal structure (Fig. 2a), that interactions within the binding pocket enable the specific recognition of the aspartate, threonine, and arginine residues of the CFTR PDZ-binding motif. These findings were corroborated by MD and binding free energy studies, where the alanine substitution mutants were found to be less energetically favourable than the WT binding motif. Interestingly, ATRL and DTAL sequences correspond to the PDZ-binding motifs of the sodium-dependent phosphate co-transporter and the H^+^ ATPase, respectively, both of which have been shown to bind to the PDZ domains of NHERF1 [34, 42–44]. As such, the PDZ-binding motif of CFTR appears to be an optimal PDZ-binding motif for interaction with the PDZ domains of NHERF1. This indicates that while the consensus motif X-S/T-X-Φ represents the minimal requirements for interaction with type I PDZ domains, residue selection at P-1 and P-3 does appear to influence the affinity of the interaction. There are several instances where competition between different PDZ domain-containing proteins has been shown to play a role in dynamic regulation of cell signalling [10, 45–47]. As such, the ability of PDZ domains to bind to both optimal and sub-optimal motifs may be key to maintaining promiscuity of the PDZ domain-motif interaction, whilst also allowing certain interactions to take precedence over others in the context of the PDZ domain interaction network.

The approach presented here demonstrates that iBox-PAK4cat crystals represent a potentially useful tool for the study of PDZ domain-motif interactions in a cellular context. The advantages of the iBox-PAK4cat system for detecting PDZ-motifs are clear: the assay is *in-cellulo*, robust, quick, with the simple read out of fluorescence, and thus is a simple assay to introduce into a cell biology laboratory. We have demonstrated here with our proof-of-principle study, that two PDZ domains can be easily be screened against a number of interacting motifs. This strategy could easily be adapted to a high-throughput format allowing for multiple PDZ domains to be screened against multiple interaction motifs, or indeed to probe other protein-protein interactions.

## Methods

### Cloning and constructs

For the generation of NHERF1 PDZ2-DTRL, cDNA for human NHERF1 PDZ2 (NHERF1 residues 150-270) was cloned into pSY5 vector [48] using standard ligation independent cloning methods [49]. The CFTR PDZ-binding motif (CFTR residues 1477-1480) was inserted downstream using Quikchange II Site-Directed Mutagenesis Kit (Agilent) according to the manufacturer’s instructions. For the generation of iBox-PAK4cat-DTRL, iBox-PAK4cat-ATRL, iBox-PAK4cat-DARL, iBox-PAK4cat-DTAL, iBox-PAK4cat-DTRA, iBox-PAK4cat-DERL, and iBox-PAK4cat-AAAA constructs, the relevant tetrapeptide motifs were inserted into iBox-PAK4cat within the pXJ40 vector [1] using the Quikchange II Site-Directed Mutagenesis Kit (Agilent) according to the manufacturer’s instructions. For the generation of GFP-PDZ1 and GFP-PDZ2 constructs NHERF1 PDZ1 (NHERF1 residues 11-120) and PDZ2 (NHERF1 residues 150–270) domains were inserted into the multiple cloning site of pXJ40-GFP [50] using XhoI and KpnI restriction sites according to standard PCR restriction cloning methods. In all cases the presence of the desired construct was confirmed through sequencing.

### Expression, purification, and crystallisation of PDZ2-DTRL

NHERF1 PDZ2-DTRL was expressed in *Escherichia coli* BL21 (DE3) (New England BioLabs). Cultures in auto-induction media (12 g/L tryptone, 24 g/L yeast extract, 9.4 g/L K_2_HPO_4_, 2.2 g/L KH_2_PO_4_, 8 mL/L glycerol 50 mM NH_4_Cl, 20 mM MgSO_4_, 5 mM Na_2_SO_4_, 0.3 % (w/v) α-lactose, 0.015 % (w/v) D-glucose) were grown at 37 °C until an OD_600_ of 0.6-0.8, following which they were grown overnight at 15 °C. Bacterial lysates were purified over HisTrap FF Ni-NTA, HiTrap Q HP, and HiLoad 16/600 Superdex 75 columns (GE healthcare) under standard conditions using a semi-automated AKTA system. Purified proteins were eluted into 50 mM Tris.HCl pH 8.0, 150 mM NaCl. SDS–PAGE and Coomassie Brilliant Blue staining assessed protein purity to be above 90 %. Crystallisation of NHERF1 PDZ2-DTRL was carried out by hanging drop method at a protein concentration of 10.5 mg/ml. Bipyramidal-shaped crystals grew in in 1.65 M ammonium sulphate and 100 mM tri-sodium citrate pH 5.6 at 15 °C.

### X-ray data collection and structure determination

Crystals were harvested into mother liquor containing 30 % (v/v) glycerol and flash-frozen in liquid nitrogen. X-ray diffraction data for NHERF1 PDZ2-DTRL were collected remotely at the Australian Synchrotron. Data were collected using an ADSC Quantum 315r CCD detector at the MX2 beamline at a wavelength of 1.0 Å. Each image covered an oscillation of 0.5 °. Images were indexed and integrated using MOSFLM [51]. The space group was determined to be I2_1_2_1_2_1_ using POINTLESS and the data was scaled using AIMLESS, both within the CCP4 suite of programs [52]. The structure was solved by molecular replacement in PHASER [53] using the structure of NHERF1 PDZ2 bound to the CXCR2 PDZ binding motif (PDB: 4Q3H) as a search model. The solvent content was determined to be 50.1 % with a Matthews coefficient (Vm) of 2.50 Å^3^/Da. This is consistent with the presence of two complexes within the asymmetric unit. Manual building of the extended PDZ domain and PDZ binding motif was carried out in Coot molecular graphics software [54]. The structure was refined to a resolution of 2.20 Å using *phenix.refine* [55], yielding a final Rwork/Rfree of 0.2360/0.2752. All figures depicting the structure were generated in PyMOL [56] unless otherwise stated.

### Transfection of cell culture

Monkey COS-7 cells were grown in high-glucose DMEM medium (Hyclone) supplemented with 10 % (v/v) bovine calf serum (Hyclone). Cells were incubated at 37 °C with 5 % CO_2_. Transfections were carried out using jetPEI transfection reagent (Polyplus transfection) according to the manufacturer’s instructions.

### Live cell imaging of crystal growth

Relevant constructs were co-transfected into COS-7 cells in 24 well black-walled glass-bottomed µ-plate (Ibidi). Immediately after transfection, images were acquired on an Eclipse Ti inverted microscope (Nikon) using a 10 x 0.3 SPlan Fluar objective, the filter sets for GFP, and a pE-300 LED (CoolLED) fluorescent light source. Imaging software NIS Elements AR.46.00.0. point visiting was used to allow five positions to be imaged for each well within the same time-course. Images were collected using a CoolSnap Monochrome camera (Photometrics) every 30 minutes for a total of 71.5 hours. GFP fluorescent images were acquired using a fixed exposure time of 200 ms. Transmitted light images were acquired using a fixed exposure time of 50 ms. Cells were maintained at 37°C and 5 % CO_2_ throughout the course of the experiment.

### Cell breakage and separation of iBox-PAK4cat crystals

Relevant constructs were co-transfected into COS-7 cells in 6-well dishes. Three days post-transfection, DMEM media was aspirated from the cells and 500 µL of hypotonic lysis buffer (50 mM Tris.HCl pH 7.4, 25 units/mL DNase I (Promega), 0.2 % (v/v) Triton-X-100), added to each well. Plates were rocked at ambient temperature until cells could be observed to detach from the plate. Lysed cells were centrifuged in a square-bottomed cuvette at ambient temperature for 5 minutes at 200 x g in a bench-top centrifuge, such that the pellet contained cell debris and the supernatant contained crystals. Supernatants were aspirated into a black-walled glass-bottomed 24 well plate (Ibidi) for imaging.

### Crystal imaging and quantification of GFP fluorescence

Harvested crystals were imaged in 24 well black-walled glass-bottomed µ-plate (Ibidi) on an MMI CellCut laser microdissection system (Olympus) using a 10 x 0.3 UPlan FLN objective, the filter sets for FITC, and a xenon mercury fluorescent light source. Images were collected using an Orca ER C4742-95-12ERG monochrome camera (Hamamatsu) using a fixed exposure time of 100 ms. Fluorescence intensity quantification was carried out using Fiji software [57]. Crystal-containing regions were selected using automatic thresholding with the Otsu algorithm and the mean grey value measured. The acquired values were plotted and subjected to two-way ANOVA with Sidak’s multiple comparisons test using Graphpad Prism 7.0 software.

### Multiple sequence alignments

Multiple sequence alignment was carried out using ClustalW2 using the default settings [58].

### Models building and Molecular Dynamics Simulations

[59]. The initial NHERF1 PDZ2-DTRL complex was edited in order to consider only the sequence portion interacting with the NHERF1 PDZ2 core domain. Subsequently the DTRL portion of the wild type peptide was modified to obtain the corresponding alanine mutants (ATRL, DARL, DTAL and DTRA). The systems were modelled using the Protein Preparation Wizard tool [60] of the Schrödinger Suite and the protonation states of H169 and H212 were evaluated and verified using the PropKa routine at pH 7.0 ± 0.2 as default. While on initial evaluation both histidines were estimated to be in HID form (neutral, δ-nitrogen protonated), on further analysis with the PropKa routine, H212 was estimated to be in the HIE state (neutral, ε-nitrogen protonated) with a pKa of 5.20, while H169 was evaluated to be present in the HIP state (+1 charged, both δ- and ε-nitrogens protonated) with a pKa value of 7.13. As a consequence, four combinations of protonation states for H169 (HID/HIP) and H212 (HID/HIE) of NHERF1 PDZ2 were evaluated. For each combination, two replicate 50 ns production runs were performed and evaluated to assess the persistence of the pKa values obtained during initial settings. Frames at defined time steps (every 5-10 ns) were selected and evaluated. All the investigated frames of WT simulations reporting HIP169 demonstrated a well-maintained HIP state as predicted in earlier phases. The root-mean square deviation (RMSD) of the Cα atoms was evaluated for 1) the whole complex, 2) the secondary structure of the PDZ2 domain, and 3) the CFTR PDZ-binding motif (Supplementary Fig.10). Generally, the RMSD value of the Cα atoms in the secondary structure was found to remain below 1.0 Å in all of the systems. Nevertheless, an improved stabilization of the peptide was observed for simulations reporting the HIP state for H169. In particular, for the HIP169/HID212 state, the RMSD value related to the Cα atoms was found to oscillate between 1.0 and 3.0 Å and the calculated binding free energies with both MM-PBSA and the SIE methods indicated that this system had much higher stability than the other systems. As such, our analyses focused on the HIP169/HID212 replicas. For simulations employing the DARL, DTAL and DTRA alanine substitution mutants, the same protonation states as described for the WT complex were employed. However, for the ATRL alanine substitution mutant, the absence of aspartate at the P-3 of the binding motif was found to significantly change the local environment so that the pKa value of H169 was decreased. As a result, for this system only the epsilon nitrogen of H169 was considered to be protonated. This procedure has been repeated to verify the pKa at fixed time steps during all simulations. MD simulations were carried out using the AMBER package, version 14. All complexes were placed in a rectangular parallelepiped water-box and solvated with a 25 Å water cap, employing the TIP3P explicit solvent model and different counterions to neutralize the systems. Simulations were conducted using the ff14SB force field at 300K, Particle mesh Ewald (PME) method for long-range electrostatic interactions and periodic boundary conditions. A cut-off of 10 Å for the non-bonded interactions was also included and the general time step for all simulations was set as 2.0 fs. The SHAKE algorithm was applied to keep all bonds involving hydrogen atoms rigid. Prior to the simulations, each system was subjected to an energy minimization using the steepest descent algorithm, followed by conjugate gradient. The temperatures of the systems were raised from 0 to 300K within 500 ps of simulation time under constant-volume periodic boundary condition. During both stages, all the Cα atoms of the complexes were kept restrained with a harmonic force constant of 10 kcal/mol⋅Å2. Multiple equilibration stages were lately performed with a constant-pressure periodic boundary condition and a gradual decrease of the harmonic force constant, until the whole system was completely relaxed. During this stage (equivalent to 3-5 ns), the pressure and the temperature of the systems were kept constant by employing the Monte Carlo barostat (with anisotropic pressure scaling) and the Langevin thermostat. Molecular dynamics productions were finally carried out with the same temperature and pressure conditions but without restraints for a singular amount of 50 ns for each peptide-protein complex. The CPPTRAJ and PTRAJ modules of AmberTools were employed to process and analyze all trajectories while UCSF Chimera [32] was utilized for visualization and preliminary inspections. RMSD plots and all images were generated using respectively Gnuplot and PyMOL [56].

### Binding free energy studies

Binding energy studies were carried out by applying the Molecular Mechanics Poisson–Boltzmann Surface Area (MM-PBSA) [61–63] method as general standard and Solvation Interaction Energies (SIE) method [11–14] in line with an approach employed in previously [15]. For the calculation of binding energies using the MM-PBSA method [61–63], the MM-PBSA implementation of the AMBER 14 package was used. The Van der Waals, electrostatics, and internal interaction were estimated by employing the SANDER module, while the MM-PBSA module was used to calculate polar energies. Dielectric constants of 1 and 80 were applied to represent the gas and water phases, respectively. MOLSURF was employed to estimate the non-polar energies, while the entropic term was deemed as approximately constant in comparison with the systems interactions. For the calculation of binding energies using the SIE method, an adapted version of the SIETRAJ tool (available at https://www2.bri.nrc.ca/ccb/pub/sietraj_main.php) was used. The SIETRAJ tool employed several scripts based on a Tcl runtime library (BRIMM) and its internal Poisson solver and surface generation methods [64–66].

### Accession codes

Atomic coordinates and structure factor files have been deposited in the Protein Data Bank under accession code 6RQR.

## Supporting information

PDB-validation-report

## Acknowledgements

We thank the University of Manchester, U.K., and the Agency for Science, Technology and Research (A*STAR), Singapore, for support. The authors thank the technical services provided by the Australian Synchrotron, part of ANSTO. All MD simulations were conducted using the Computational Shared Facility at The University of Manchester. Imaging of the iBox-PAK4cat crystals was carried out at the Bioimaging Facility at the University of Manchester. We wish to thank Dr Mattia Mori (University of Siena, Italy), Dr Giulio Poli (University of Pisa, Italy), Dr Hao Fan (BII, Singapore), Dr Peter March (University of Manchester, U.K.), Dr Bo Xue (NUS, Singapore) Dr Yohendran Baskaran (IMCB, Singapore), Dr Moses Kakanga (IMCB, Singapore), Dr Ed Manser (IMCB, Singapore), and the IT staff of The University of Manchester for their support.

## Author contributions

R.C.R., R.C.F., and E.R.M. conceived the study. R.C.R., R.C.F., E.R.M., and A.B. wrote the paper. E.R.M. designed and generated cDNA plasmid constructs and carried out all *in-vitro* and *in-vivo* crystallisation experiments. R.C.R. and E.R.M. carried out X-ray data collection and structure refinement. A.B. carried out MD simulations and binding energy studies.

