## Supplementary tables and figures for "*In-vivo* crystals reveal protein interactions"

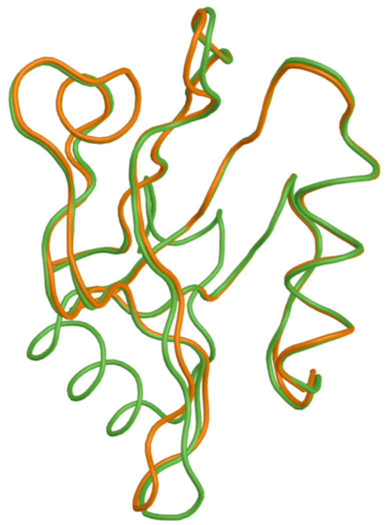

**Supplementary Figure 1:** Extended PDZ2 domain of NHERF1 obtained in this study (green) is overlaid with the crystal structure of the canonical domain (PDB: 2OZF) (orange).

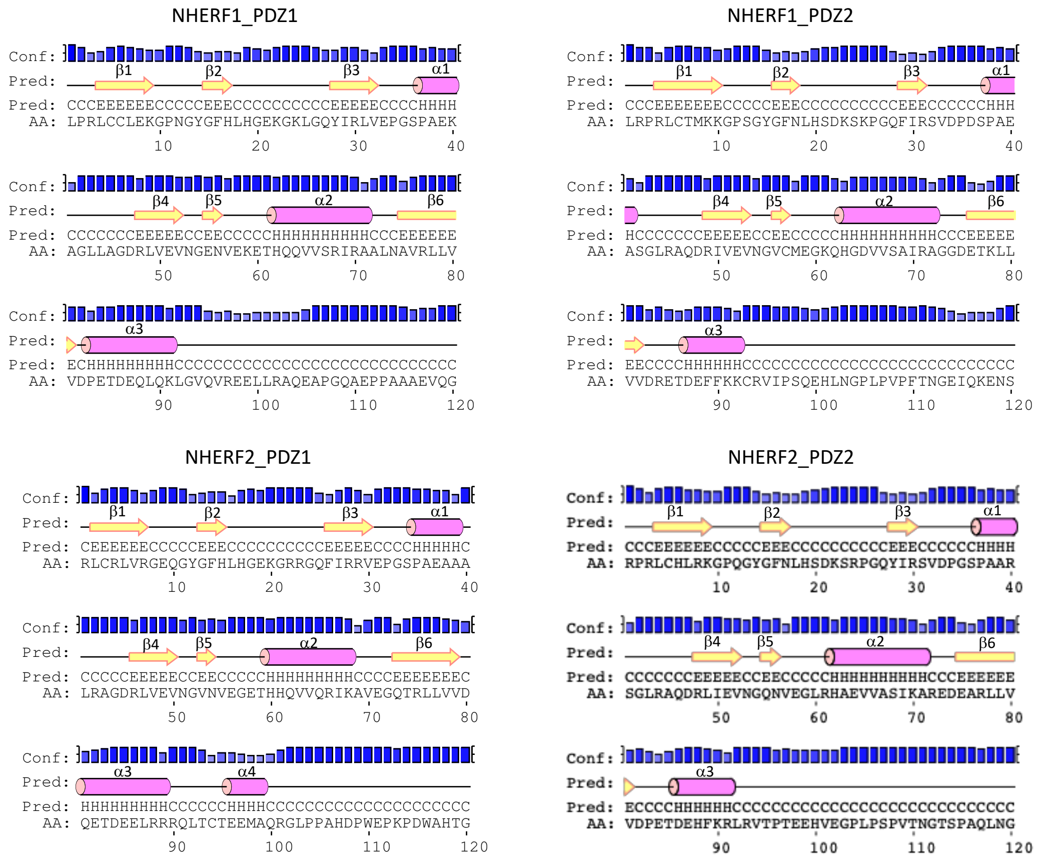

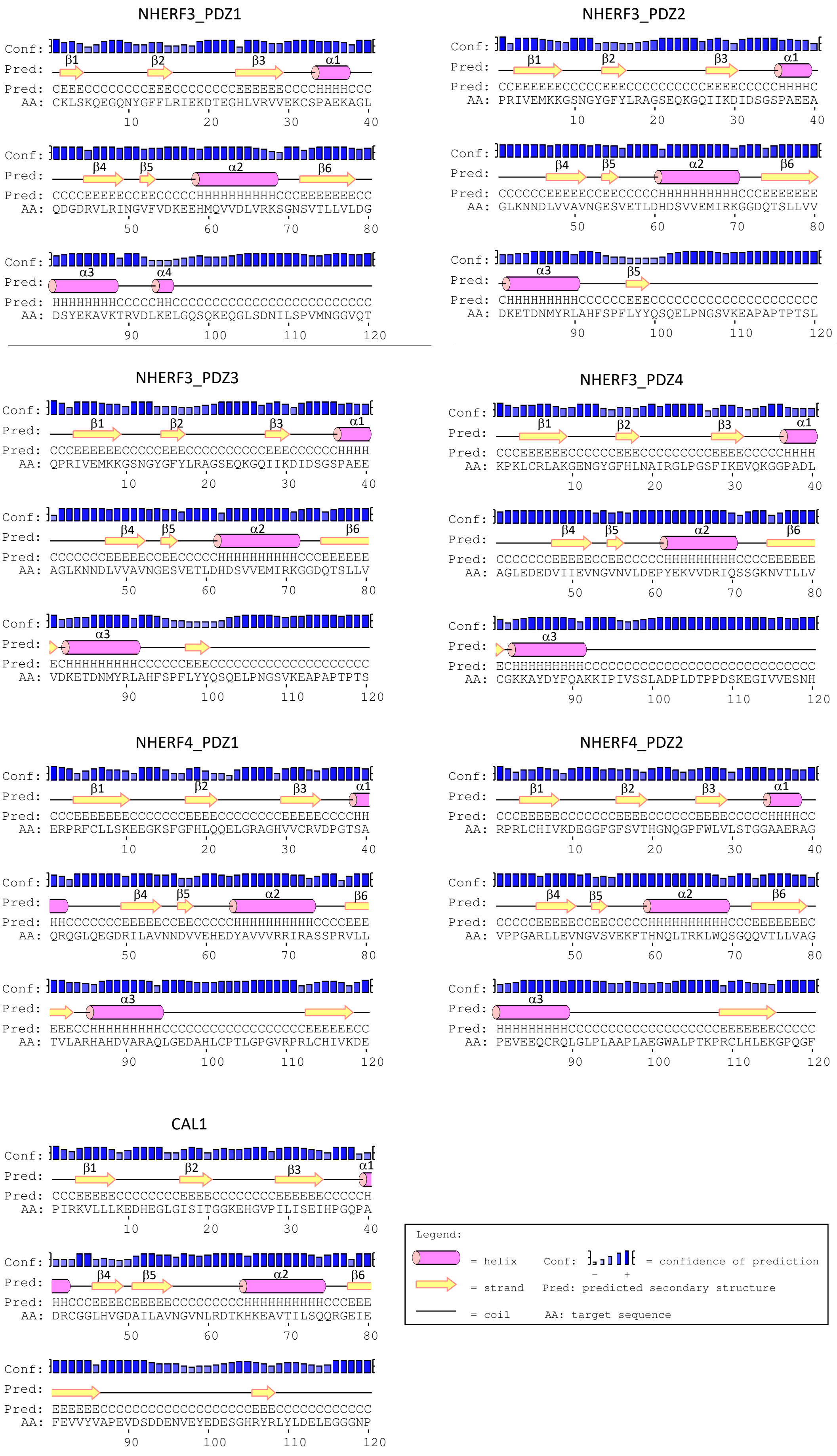
NHERF3_PDZ1
Conf:
Pred: β1 β2 β3 α1
Pred: CEEECCCCCCCCEEECCCCCCCCEEEEEEECCCCHHHHCCC
AA: CKLSKQEGQNYGFFLRIEKDTEGHLVRVVEKCSPAEKAGL
10 20 30 40
Conf:
Pred: β4 β5 α2 β6
Pred: CCCCEEEEEECCEECCCCCHHHHHHHHHHHCCCEEEEEEEECC
AA: QDGDRVLRINGVFVDKEEHMQVVDLVRKSGNSVTLLVLDG
50 60 70 80
Conf:
Pred: α3 α4
Pred: HHHHHHHHCCCCCHHCCCCCCCCCCCCCCCCCCCCCCCCC
AA: DSYEKAVKTRVDLKELGQSQKEQGLSDNILSPVMNGGVQT
90 100 110 120
NHERF3_PDZ2
Conf:
Pred: β1 β2 β3 α1
Pred: CCEEEEEEECCCCCEEECCCCCCCCCCCEEEECCCCCHHHHC
AA: PRIVEMKKGSNGYGFYLRAGSEQKGQIIKDIDSGSPAEEA
10 20 30 40
Conf:
Pred: β4 β5 α2 β6
Pred: CCCCCCEEEEEECCEECCCCCHHHHHHHHHHCCCEEEEEEEE
AA: GLKNNDLVVAVNGESVETLDHDSVVEMIRKGGDQTSLLVV
50 60 70 80
Conf:
Pred: α3 β5
Pred: CHHHHHHHHHCCCCCCEEECCCCCCCCCCCCCCCCCCCCC
AA: DKETDNMYRLAHFSPFLYYQSQELPNGSVKEAPAPTPTSL
90 100 110 120
NHERF3_PDZ3
Conf:
Pred: β1 β2 β3 α1
Pred: CCCEEEEEEECCCCCEEECCCCCCCCCCCEEECCCCCCHHHH
AA: QPRIVEMKKGSNGYGFYLRAGSEQKGQIIKDIDSGSPAEE
10 20 30 40
Conf:
Pred: β4 β5 α2 β6
Pred: CCCCCCCEEEEEECCEECCCCCHHHHHHHHHHCCCEEEEEEE
AA: AGLKNNDLVVAVNGESVETLDHDSVVEMIRKGGDQTSLLV
50 60 70 80
Conf:
Pred: α3
Pred: ECHHHHHHHHHCCCCCCEEECCCCCCCCCCCCCCCCCCCC
AA: VDKETDNMYRLAHFSPFLYYQSQELPNGSVKEAPAPTPTS
90 100 110 120
NHERF3_PDZ4
Conf:
Pred: β1 β2 β3 α1
Pred: CCCEEEEEEECCCCCCEEECCCCCCCCCCEEEECCCCCHHHH
AA: KPKLCRLAKGENGYGFHLNAIRGLPGSFIKEVQKGGPADL
10 20 30 40
Conf:
Pred: β4 β5 α2 β6
Pred: CCCCCCCEEEEEECCEECCCCCHHHHHHHHHCCCCEEEEEEE
AA: AGLEDEDVIIEVNGVNVLDEPYEKVVDRIQSSGKNVTLLV
50 60 70 80
Conf:
Pred: α3
Pred: ECHHHHHHHHHCCCCCCCCCCCCCCCCCCCCCCCCCCCCC
AA: CGKKAYDYFQAKKIPIVSSLADPLDTPPDSKEGIVVESNH
90 100 110 120
NHERF4_PDZ1
Conf:
Pred: β1 β2 β3 α1
Pred: CCCEEEEEEEECCCCCCCEEEECCCCCCCCEEEEEECCCCHH
AA: ERPRFCLLSKEEGKSFGFHLQQELGRAGHVVCRVDPGTSA
10 20 30 40
Conf:
Pred: β4 β5 α2 β6
Pred: HHCCCCCCCEEEEEECCEECCCCCHHHHHHHHHHCCCCEEE
AA: QRQGLQEGDRILAVNNDVVEHEDYAVVVRRIRASSPRVLL
50 60 70 80
Conf:
Pred: α3
Pred: EEECCHHHHHHHHHCCCCCCCCCCCCCCCCCCEEEEEEECC
AA: TVLARHAHDVARAQLGEDAHLCPTLGPGVRPRLCHIVKDE
90 100 110 120
NHERF4_PDZ2
Conf:
Pred: β1 β2 β3 α1
Pred: CCCEEEEEECCCCCCCEEEEECCCCCCEEEECCCCCHHHHCC
AA: RPRLCHIVKDEGGFGFSVTHGNQGPFWLVLSTGGAAERAG
10 20 30 40
Conf:
Pred: β4 β5 α2 β6
Pred: CCCCCEEEEECCEECCCCCHHHHHHHHHHCCCEEEEEEEEC
AA: VPPGARLLEVNGVSVEKFTHNQLTRKLWQSGQQVTLLVAG
50 60 70 80
Conf:
Pred: α3
Pred: HHHHHHHHHCCCCCCCCCCCCCCCCCCEEEEEEEECCCCC
AA: PEVEEQCRQLGLPLAAPLAEGWALPTKPRCLHLEKGPQGF
90 100 110 120
CAL1
Conf:
Pred: β1 β2 β3 α1
Pred: CCCEEEEEECCCCCCCCCEEEEECCCCCCCCCEEEEEEECCCCCH
AA: PIRKVLLLKEDHEGLGISITGGKEHGVPILISEIHPGQPA
10 20 30 40
Conf:
Pred: β4 β5 α2 β6
Pred: HHCCCEEEECEEEEECCCCCCCCCHHHHHHHHHHCCCEEE
AA: DRCGGLHVGDAILAVNGVNLRDTKHKEAVTILSQQRGEIE
50 60 70 80
Conf:
Pred:
Pred: EEEEEECCCCCCCCCCCCCCCCCCCCEEECCCCCCCCCCCCC
AA: FEVVYVAPEVDSDDENVEYEDESGHRYRLYLDELEGGGNP
90 100 110 120
Legend:
= helix
Conf: = confidence of prediction
− +
= strand
Pred: predicted secondary structure
= coil
AA: target sequence

**Supplementary Figure 2:** Secondary structure prediction for the PDZ domains of the NHERF family. A third helix ($\alpha$3) after the $\beta$6 strand is predicted for several PDZ domains within the NHERF family, but not for those outside such as CAL1. Note: A fourth helix ($\alpha$4) is predicted only for NHERF2 PDZ1 and NHERF3 PDZ1, but is not predicted to be present in either NHERF1 PDZ1 or PDZ2. Secondary structure predictions were carried out using the PSIPRED server v3.3 (available at http://bioinf.cs.ucl.ac.uk/psipred) using the default settings.

```
Human      ISPSDRVKLFP--HRNSSKCKSKPQIA----ALKEETEEEVQDTRL 1480
Frog       --HSDRLKLFPLHRRNSSKRKSRPQIS----ALQEETEEEVQDTRL 1485
Rat        LSSSEKMKLFH--GRHSSKQKPRTQIT----AVKEETEEEVQETRL 1476
Mouse      ISSSEKMRFFQ--GRHSSKHKPRTQIT----ALKEETEEEVQETRL 1476
Zebrafish  ----ERLKLFP--RRNSSMRTPQSKLSSVTQTLQEEAEDNIQDTRL 1485
Dog        IGPPERPGLLP--HRLSSRQRSPSRIA----ALKEETEDEVQDTRL 1483
Horse      ISSSDPLKLFP--HRNSSKHKSRSKIA----ALQEETEEEVQETRL 1481
Bovine     ISPADRLKLLP--HRNSSRQRSRSNIA----ALKEETEEEVQETKL 1481
Rabbit     ISSSDRAKLFP--HRNSSKHKSRPQIT----ALKEEAEEEVQGTRL 1476
Sheep      ISPADRLKLLP--HRNSSRQRSRANIA----ALKEETEEEVQETKL 1481
               :   ::    * **      .::    :::**:*:::* *:*
```

**Supplementary Figure 3:** Cross-species sequence alignment of the CFTR C-terminus. Last 40 residues of CFTR were aligned using ClustalW2 with the default settings.

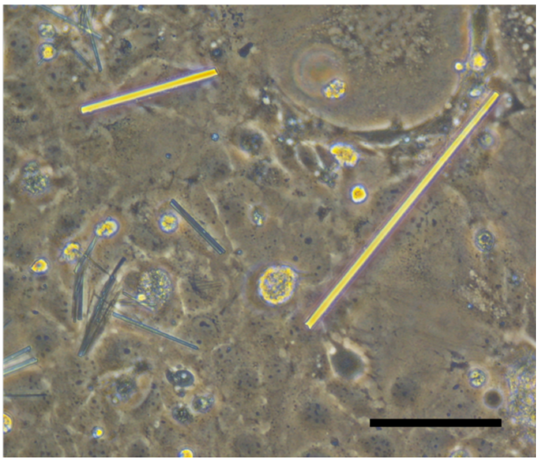

**Supplementary Figure 4:** *In-cellulo* crystals formed by iBox-PDZ2-PAK4cat fusion construct. Crystals were imaged four days post-transfection in COS7 cells. Scale bar indicates 100 µm.

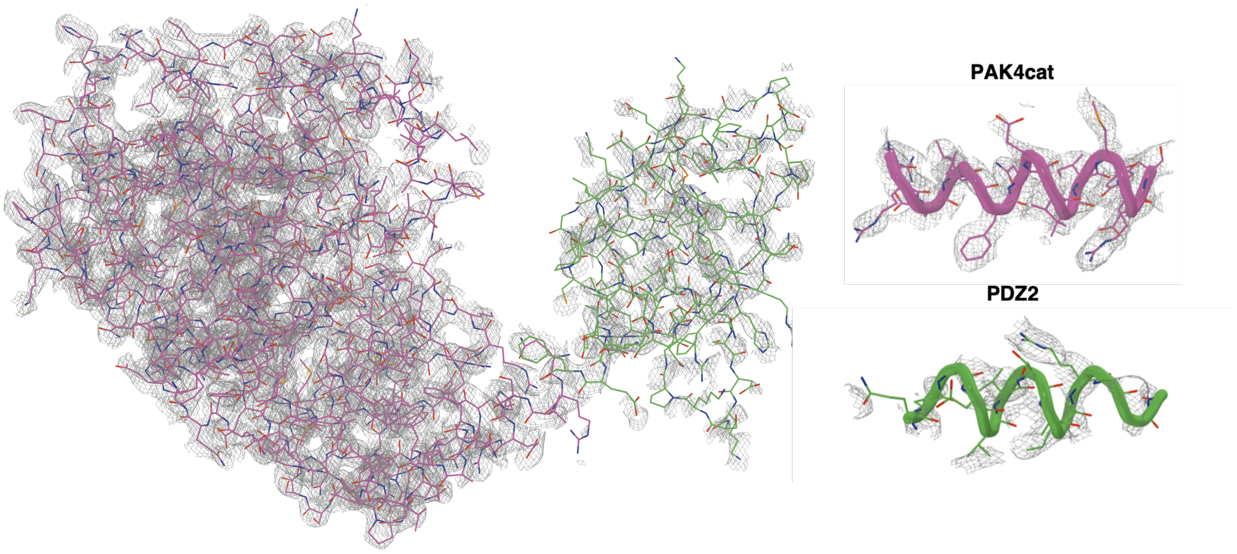


**Supplementary Figure 5:** Unrefined electron density map obtained for the iBox-PDZ2-PAK4cat fusion construct. OMIT mesh map is contoured at 1.0 sigma and within 1.5 Å of the respective atoms. Left, iBox-PAK4cat is shown in magenta and PDZ2 in green. Right, comparison of electron density observed for typical helices in PAK4cat and PDZ.

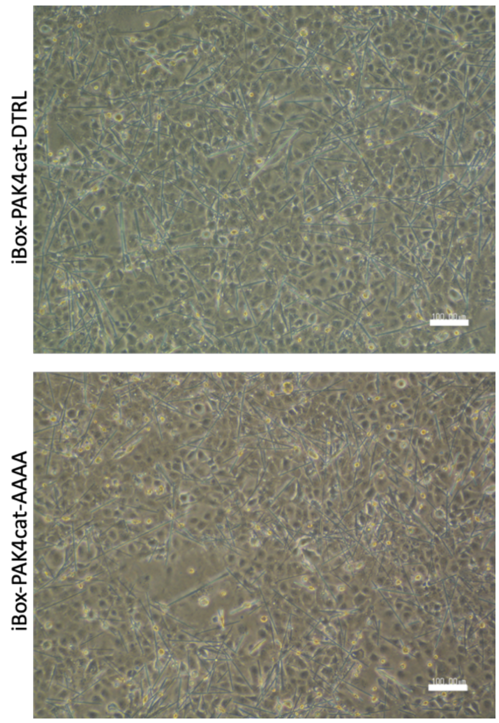


**Supplementary Figure 6:** Light images of COS-7 cells co-transfected with GFP-PDZ1 and either iBox-PAK4cat-DTRL or iBox-PAK4cat-AAAA as indicated. Both iBox-PAK4cat construct variants appear to be similar in terms of crystal generation proficiency.

**GFP-PDZ1**

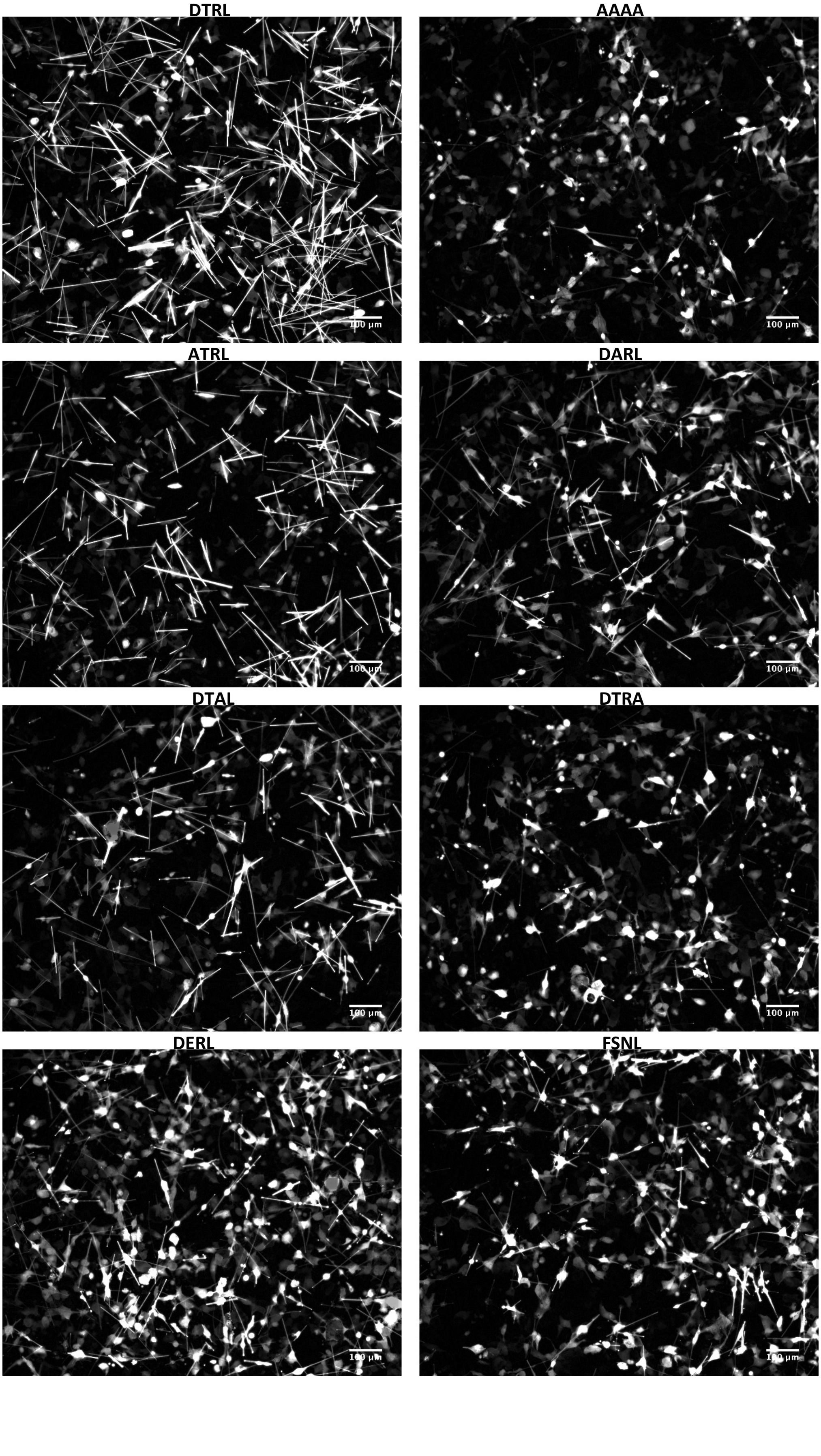

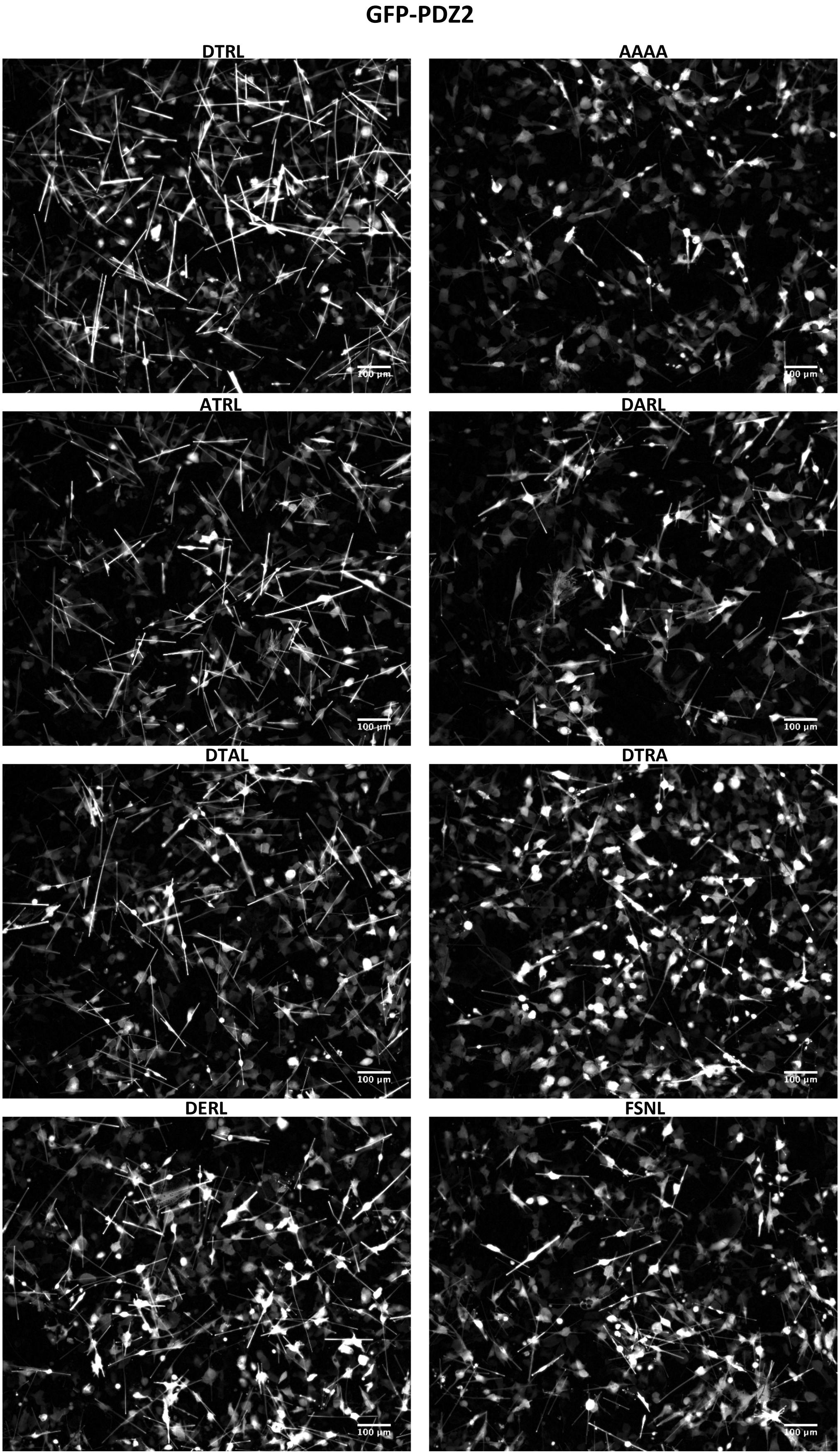


**Supplementary Figure 7:** Representative GFP fluorescent images of COS-7 cells co-transfected with GFP-PDZ and iBox-PAK4-DTRL construct variants as indicated.

**GFP-PDZ1**

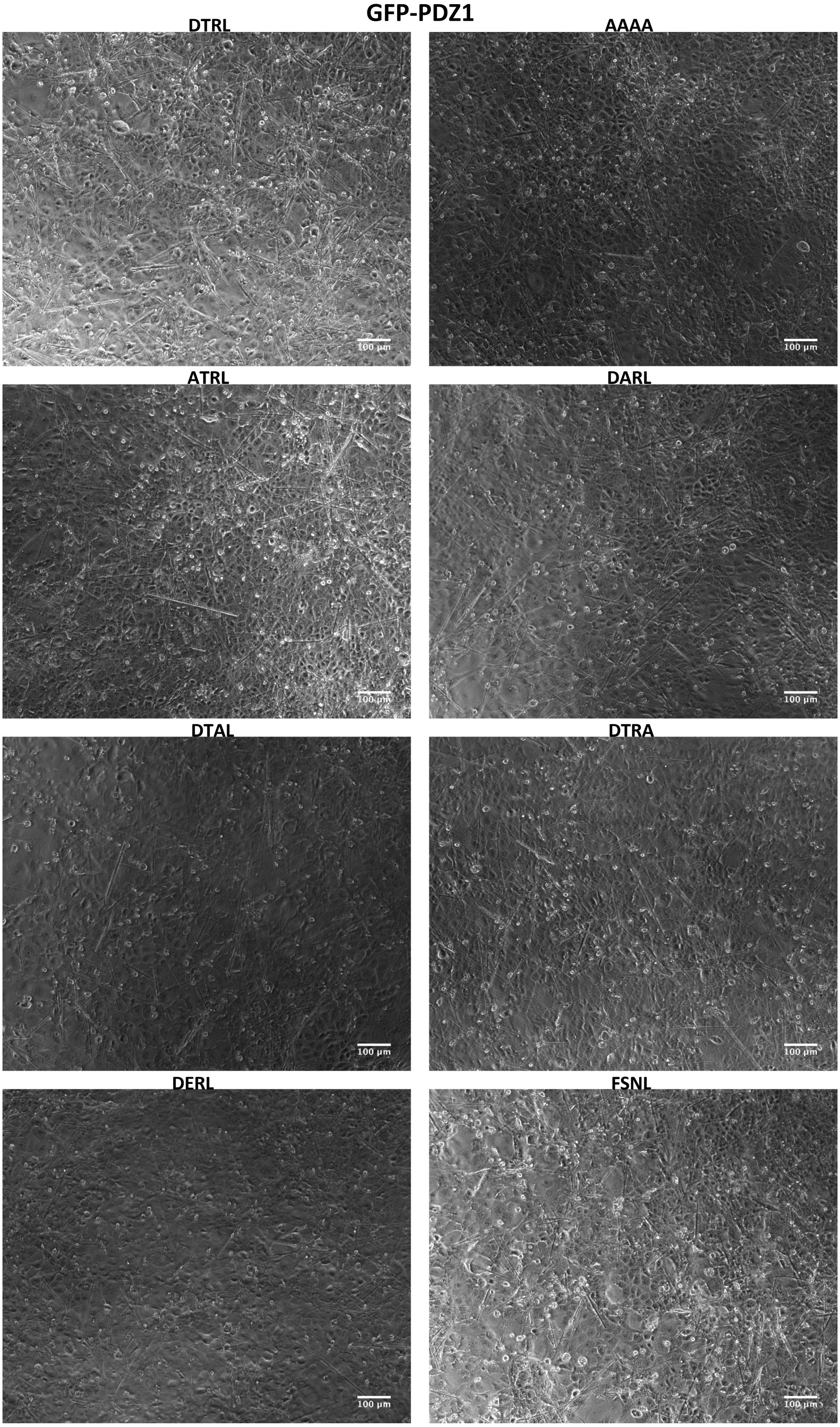

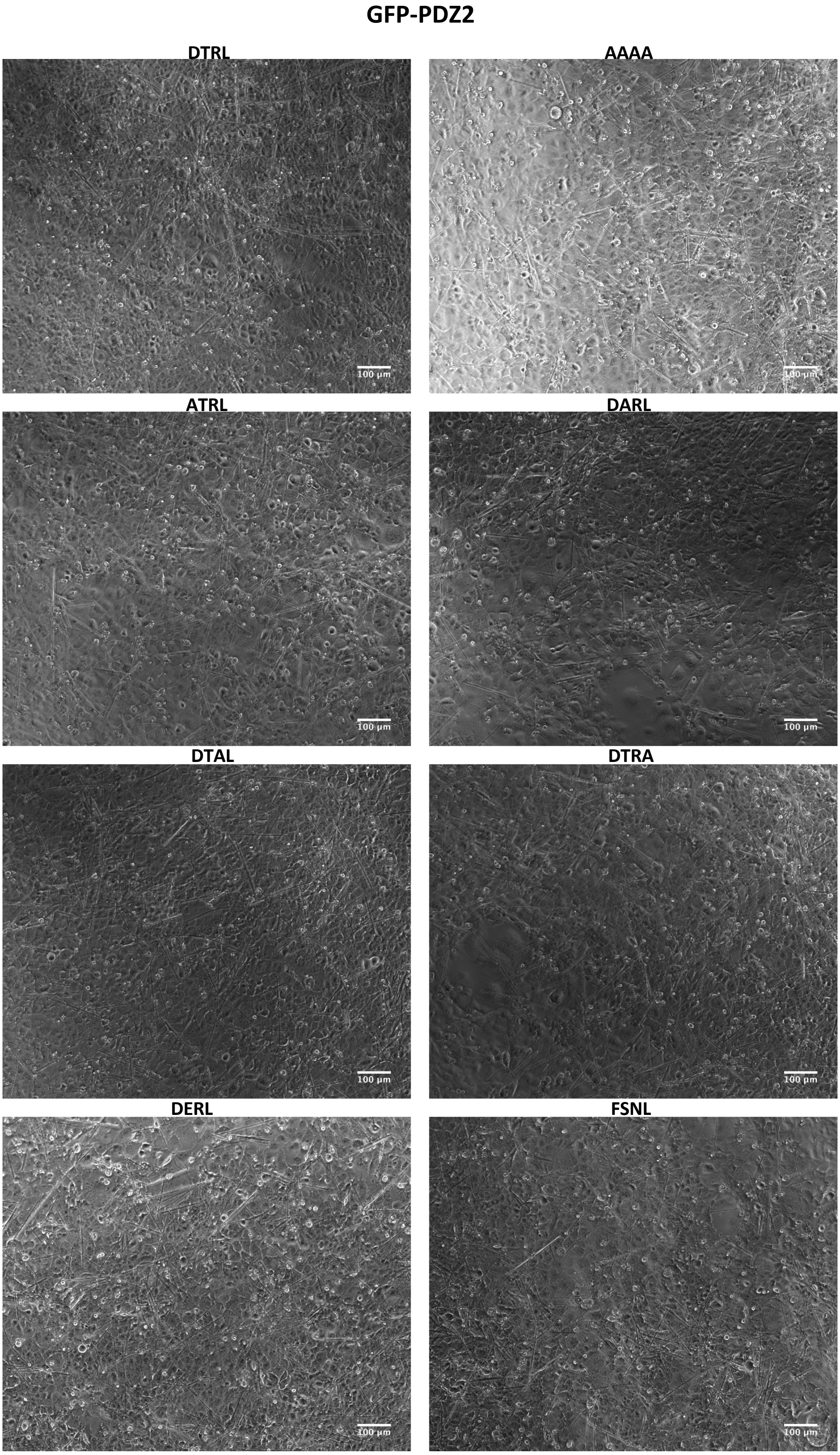


**Supplementary Figure 8:** Corresponding light images for Supplementary Figure 7.

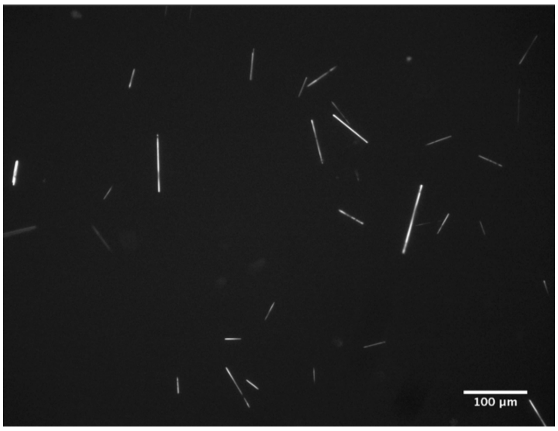


**Supplementary Figure 9:** GFP fluorescent image of crystals obtained from lysed cells following co-transfection with GFP-PDZ1 and iBox-PAK4cat-DTRL constructs

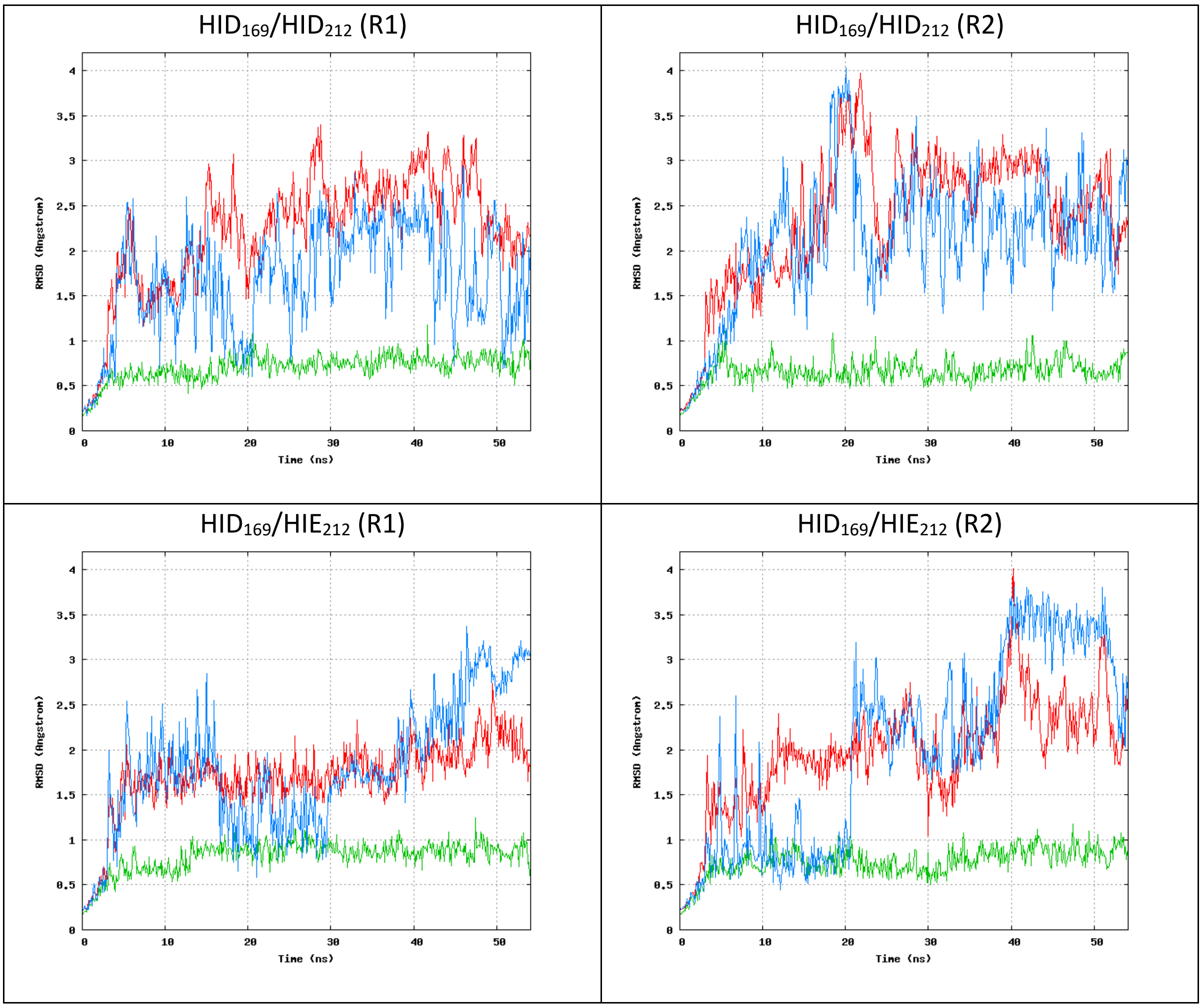

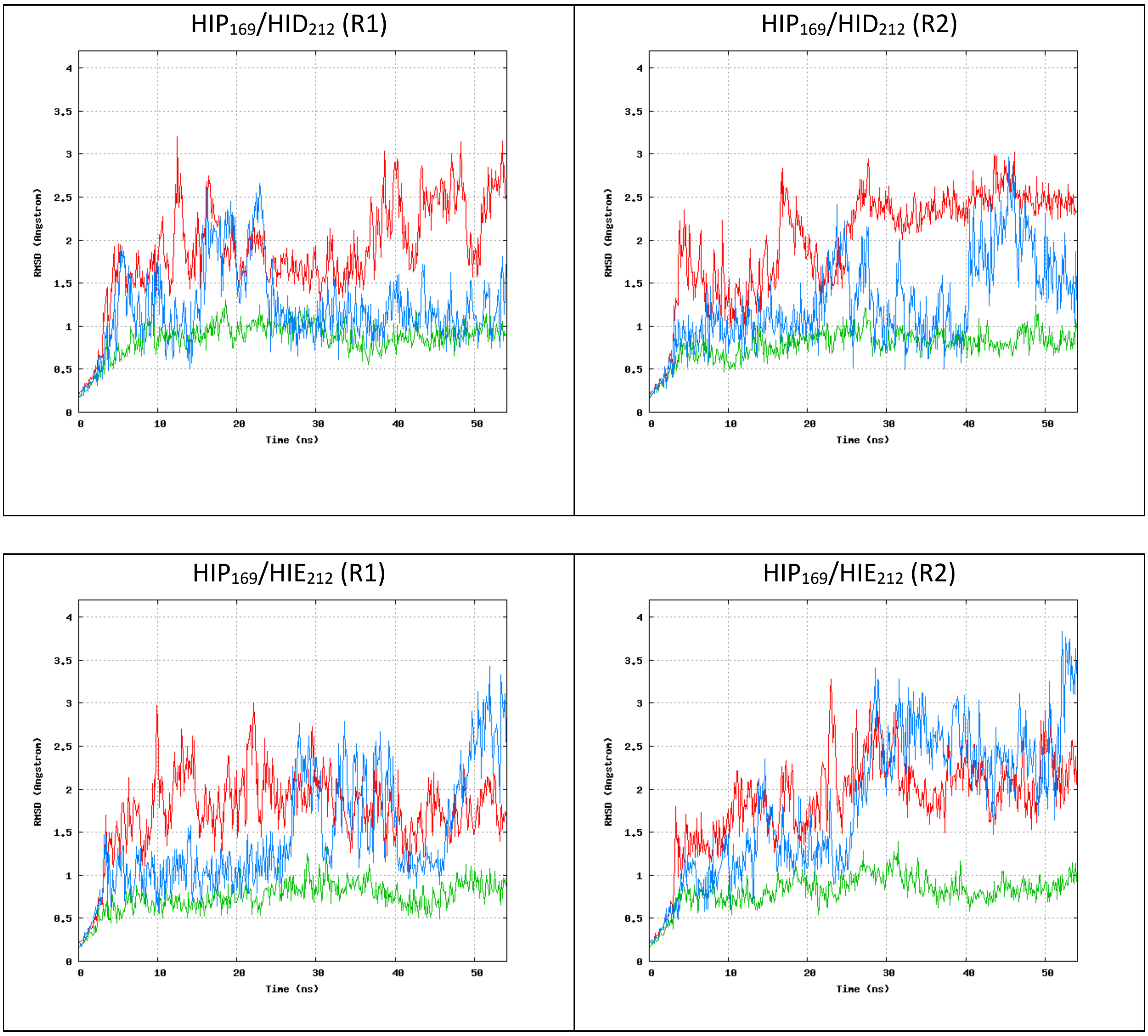


**Supplementary Figure 10:** Root-mean-square-deviation (RMSD) of NHERF1 PDZ2 in complex with the CFTR PDZ-binding motif with different protonation states for H169 and H212. HIP (+1 charged, both δ- and ε-nitrogens protonated), HID (neutral, δ-nitrogen protonated), and HIE (neutral, ε-nitrogen protonated). R = replicate. RMSD for all Cα atoms is shown in red, the RMSD related to Cα atoms of the secondary structure elements of the complex is shown in green, and the RMSD of Cα atoms of the peptide is shown in blue.

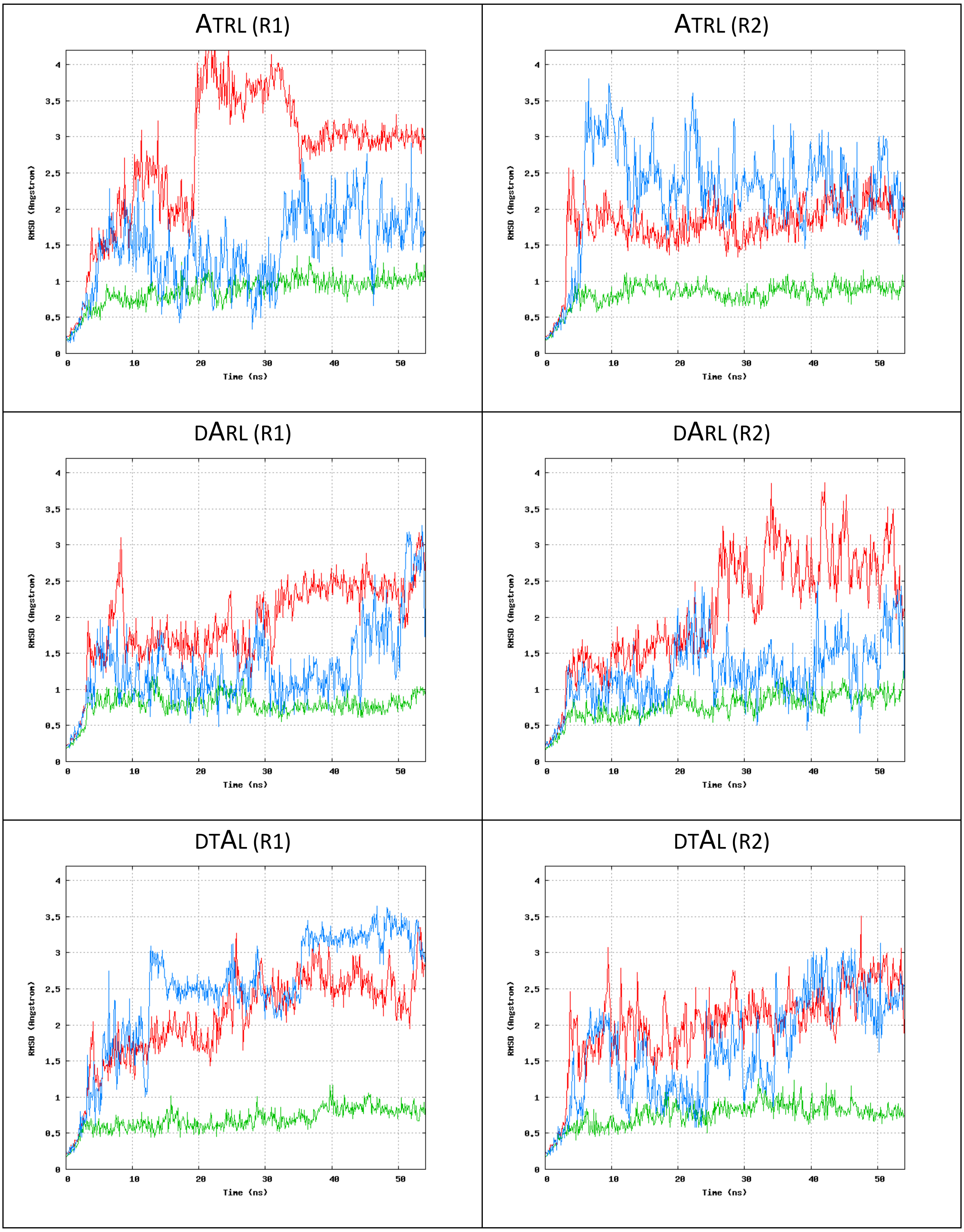

ATRL (R1)
ATRL (R2)
DARL (R1)
DARL (R2)
DTAL (R1)
DTAL (R2)
RMSD (Angstrom)
Time (ns)

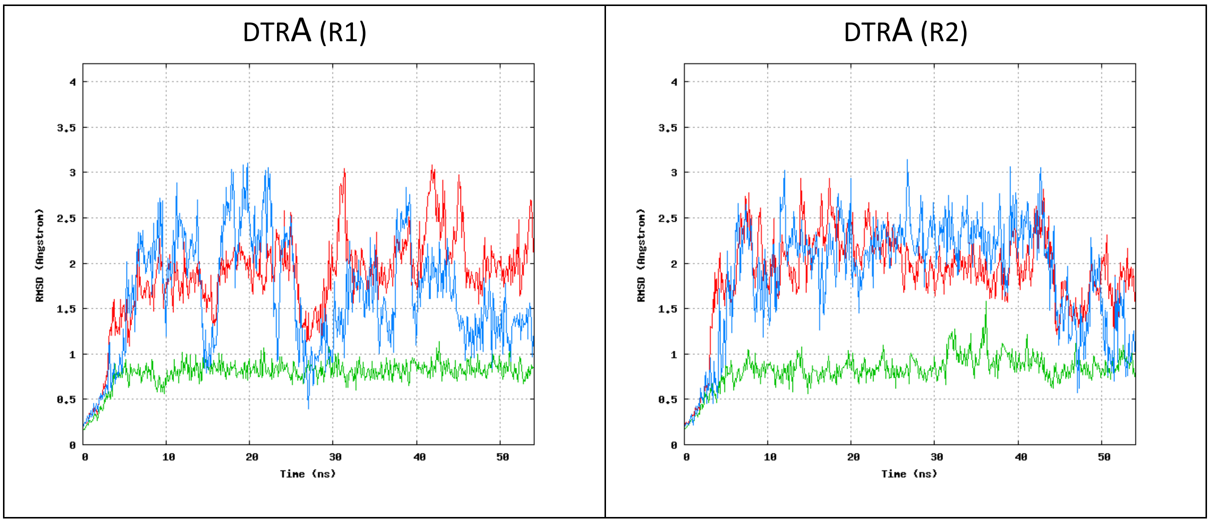


**Supplementary Figure 11:** Root-mean-square-deviation (RMSD) of NHERF1 PDZ2 in complex with the CFTR PDZ-binding motif alanine substitution mutants as indicated. R = replicate. RMSD for all Cα atoms is shown in red, the RMSD related to Cα atoms of the secondary structure elements of the complex is shown in green, and the RMSD of Cα atoms of the peptide is shown in blue.

| | |
|---|---|
| **Space group** | $P6_3$ |
| **Cell dimensions** | |
| (a, b, c) (Å) | 143.1, 143.1, 61.8 |
| (α, β, γ) (°) | 90.0, 90.0, 120.0 |
| **Resolution range (Å)** | 19.9-3.15 (3.21-3.15) |
| **Unique reflections** | 13276 (633) |
| **Completeness (%)** | 99.9 (99.4) |
| **Multiplicity** | 3.6 (3.1) |
| **$R_{merge}$** | 18.8 (57.6) |
| **I/σ** | 6.3 (2.0) |

**Supplementary Table 1:** Diffraction statistics for the iBox-PDZ2-PAK4cat fusion crystal.

**Supplementary Movie 1:** Fluorescent time-lapse comparison of COS-7 cells co-transfected with GFP-PDZ2 and iBox-PAK4cat-DTRL, left, or iBox-PAK4catDTRA, right.
